# *In Vitro* Modulator Responsiveness of 655 *CFTR* Variants Found in People With CF

**DOI:** 10.1101/2023.07.07.548159

**Authors:** Hermann Bihler, Andrey Sivachenko, Linda Millen, Priyanka Bhatt, Amita Thakerar Patel, Justin Chin, Violaine Bailey, Isaac Musisi, André LaPan, Normand E. Allaire, Joshua Conte, Noah R. Simon, Amalia S. Magaret, Karen S. Raraigh, Garry R. Cutting, William R. Skach, Robert J. Bridges, Phil J. Thomas, Martin Mense

**Affiliations:** CFFT Lab, Cystic Fibrosis Foundation, Lexington, MA 02421, USA; University of Texas Southwestern Medical Center, Dallas, TX 75390, USA; Rosalind Franklin University Medical School, Chicago, IL 60064, USA; University of Washington, Seattle, WA 98195-9300, USA; Johns Hopkins University School of Medicine, Baltimore, MD 21205-2196, USA; Cystic Fibrosis Foundation, Bethesda, MD 20814, USA

**Keywords:** Cystic Fibrosis, CFTR, genetic variants, theratyping, CFTR modulator, cell-based

## Abstract

**Background:** In 2017, the US Food and Drug Administration initiated expansion of drug labels for the treatment of cystic fibrosis (CF) to include CF transmembrane conductance regulator (CFTR) gene variants based on *in vitro* functional studies. This study aims to identify *CFTR* variants that result in increased chloride (Cl^-^) transport function by the CFTR protein after treatment with the CFTR-modulator combination elexacaftor/tezacaftor/ivacaftor (ELX/TEZ/IVA). These data may benefit people with CF (pwCF) who are not currently eligible for modulator therapies.

**Methods:** Plasmid DNA encoding 655 CFTR variants and wild-type (WT) *CFTR* were transfected into Fisher Rat Thyroid cells that do not natively express CFTR. After 24 hours of incubation with control or TEZ and ELX, and acute addition of IVA, CFTR function was assessed using the transepithelial current clamp conductance assay. Each variant’s baseline activity, responsiveness to IVA alone, and responsiveness to the TEZ/ELX/IVA combination were measured in three different laboratories. Western blots were conducted to evaluate CFTR protein maturation and complement the functional data.

**Results and Conclusions:** 253 variants not currently approved for CFTR modulator therapy showed low baseline activity (<10% of normal CFTR Cl^-^ transport activity). For 152 of these variants, treatment with ELX/TEZ/IVA improved the Cl^-^ transport activity by ≥10% of normal CFTR function, which is suggestive of clinical benefit. ELX/TEZ/IVA increased CFTR function by ≥10 percentage points for an additional 140 unapproved variants with ≥10% but <50% of normal CFTR function at baseline. These findings significantly expand the number of rare CFTR variants for which ELX/TEZ/IVA treatment should result in clinical benefit.

## Introduction

Cystic fibrosis (CF) is a progressive autosomal recessive condition affecting over 40.000 people in the United Stated and over 100.000 people worldwide. CF is caused by specific variants^1^ of the *CFTR* gene that result, through various mechanisms, in a dysfunctional CFTR protein. More than 4000 *CFTR* variants have been reported, including over 1000 exonic variants (www.genet.sickkids.on.ca, Ideozu+, 2023 [1]), but not all cause disease. To date, 1654 variants have been reported in pwCF to the **C**linical and **F**unctional **Tr**anslation of CFTR (CFTR2) project (https://cftr2.org). The pathogenicity of 804 variants has been determined and almost all variants observed in at least three pwCF enrolled in CFTR2 have been evaluated. However, most of the remaining 850 uncharacterized variants in CFTR2 are rare, occurring in only one or few pwCF worldwide. This ultra-low prevalence creates three challenges: 1) lack of associated clinical data to assess disease-liability, 2) substantial resources needed to assess functional effect and 3) insufficient numbers of pwCF harboring these rare variants to conduct traditional clinical studies. In recognition of the latter issue, in 2017, the US Food and Drug Administration (FDA) first allowed the use of well-controlled *in vitro* assays to generate CFTR variant-specific drug responsiveness data for FDA label expansion, paving a way for people with rare CF-causing variants to gain access to modulator therapies (Durmowicz AG+, 2018 [2]). The four FDA-approved CF medicines by Vertex Pharmaceuticals, Kalydeco®, Symdeko® (a.k.a. Symkevi®), and Trikafta® (a.k.a. Kaftrio®) have seen label expansions based on *in vitro* data, leading to a total of 183 approved variants across these three CFTR modulator therapies. Trikafta®, the most widely used CFTR-targeted medicine, is now approved in the US for 178 variants in pwCF age 2 years and older (FDA, 2021rev [3]).

When in 2020 the number of modulator therapy-approved CFTR variants increased to 183, an additional >600 pwCF in the United States (up to 2% of pwCF living in the US) became eligible for modulator therapy. The approval of treatment for rare CF-causing variants based on *in vitro* data provides a unique opportunity to extend the benefit of these therapies to people with *CFTR* genotypes that cannot be assessed in traditional clinical trials. It is the hope that eventually such data will not only be used in the US but worldwide to promote access and/or approve treatment for all people that could benefit from CFTR modulator therapies.

The purpose of this study is to characterize CFTR variants for their responsiveness to CFTR modulators, a process that has been named theratyping (Cutting, 2015 [4]; Veit+, 2016 [5]), and to thereby identify additional CF-causing variants that could be treated with modulator therapy. To this end, 655 *CFTR* variants (comprising one or several exonic variants) identified as known or putatively disease-causing were selected from the CFTR2 database. The variants consist of 590 missense variants, 28 in-frame indels, and 33 complex alleles with multiple variants. Also included were four variants not expected to produce CFTR protein to serve as negative controls (c.830G>A [p.Trp277Ter; legacy: W277X], c.2443G>T [p.Glu815Ter; legacy: E815X], c.2496C>A [p.Cys832Ter; legacy: C832X], and c.1360_1387del [p.Leu454AlafsX6; legacy: 1491_1500del]). To enable screening large numbers of CFTR variants, a cDNA-based, scalable assay was developed, suitable to theratyping missense and small in-frame insertion or deletion (indel) variants. Of note, this approach is not compatible with the assessment of promoter or intronic *CFTR* variants, nor exonic variants that cause aberrant mRNA splicing nor nonsense variants resulting in truncated, often non-expressed CFTR protein. Plasmids were individually generated for CMV promoter-driven expression of wild-type (WT) CFTR and all 655 variants. Plasmids were transiently transfected into unmodified Fisher rat thyroid (FRT) cells, which do not natively express CFTR. Electrophysiological assays measuring CFTR Cl^-^ conductance were used to determine the forskolin-induced variant baseline CFTR transport activity, which could also be considered as baseline residual activity. Also in the electrophysiological assay, responsiveness was assessed to ivacaftor (IVA) alone and to the triple modulator combination of tezacaftor (TEZ), elexacaftor (ELX), and IVA, the active ingredients of the Vertex medicine Trikafta®.

To assess reproducibility, this study was carried out in three separate laboratories: Rosalind Franklin University of Medical Sciences in North Chicago, IL, University of Texas Southwestern Medical Center, Dallas, TX, and the CF Foundation Therapeutics Lab, Lexington, MA, henceforth referred to as RF, UT, and CF, respectively. Of the 651 *CFTR* variants in this study that are predicted to yield full-length protein, 473 are not currently approved for CFTR modulator therapies, and most had not previously been assessed for modulator responsiveness. Variants where treatment with the combination ELX/TEZ/IVA showed increased Cl^-^ transport function by at least 10% of WT CFTR function were categorized as responsive, consistent with the threshold applied for the FDA label expansions based on other *in vitro* data. There is reasonable evidence based on existing clinical and correlated *in vitro* data that achieving this level of response in a functional assay is predictive of clinical benefit across a cohort of pwCF (Durmowicz AG+, 2018 [2]). While only the drug sponsor can seek FDA-approval for additional variants, this study hopes to provide guidance to the drug sponsor, pwCF and their treating physicians regarding currently unapproved *CFTR* variant genotypes that may benefit from treatment with CFTR modulators.

## Methods

### CFTR variants

WT CFTR cDNA used in this study contained full length CDS derived from RefSeq NM000492 with the inclusion of 61 bp of the 5’ UTR immediately upstream of the CDS and the following additional changes. The CDS contained c.1408G>A variant (p.Val470Met; legacy: V470M), as well as four silent substitutions (synonymous changes): c.798T>C (p.Ile266=), c.801A>G (p.Glu267=), c.804T>C (p.Asn268=), and c.4437G>A (p.Arg1479=). This cDNA was cloned into the pEZT-BM vector (Morales-Perez CL+, 2016 [6]) using the Xho1 and Kpn1 restriction sites and was driven by CMV promoter 576bp upstream of the retained CFTR 5’ UTR. The plasmid coding for WT CFTR served as the template for all additional variants (for a complete list of variants see **Table 2** or **Table S1**) which were generated and produced to scale by Quintara Biosciences (USA) or by Genewiz (now Azenta Life Sciences, USA). pDNA was received at maxiprep scale at CF, dispensed into a master and three daughter plates. CFTR plasmids were randomly arrayed into the 96-well plates and assay operators were blinded with regards to plasmid variant ID on any of the plates. The daughter plates were distributed to the participating labs and served the experiments carried out at the three contributing labs.

To rule out any detrimental plasmid sequence defects, for final quality assurance the CFFT Lab subjected all its plasmids from the daughter plate to next generation sequencing (NGS): DNA from plasmid constructs was quantitated using Qubit dsDNA HS Assay Kit (Q33230, Thermo Fisher Scientific, USA) as per manufacturer instructions. DNA was tagmented using Nextera XT DNA library preparation kit (FC-131-1096, Illumina, USA) and dual-indexed with Nextera XT Index Kit V2 Sets A-D (FC-131-2001, FC-131-2002, FC-131-2003, FC-131-2004, Illumina, USA) according to the manufacturer’s recommendations, except in 50% reaction volumes. Fragment size distributions of tagmented and dual indexed libraries were analyzed by capillary gel electrophoresis using a Bioanalyzer High Sensitivity DNA kit (50067-4626, Agilent, USA). Up to 384 high quality libraries with size distribution between 250-1000bp were pooled using equal molar quantities. Pooled libraries were quantified using the Kapa library quantification kit (KK4824, Roche, USA). 4 nM library pools were diluted and denatured to 1.3 pM and dual index sequenced on an Illumina Miniseq using a Miniseq Mid Output 300 cycle Reagent kit (FC-420-1004, Illumina, USA). High quality Miniseq runs produced > 2.0Gb, > 85% bases Q30, > 14 million paired end reads.

Variant plasmids from 96-well daughter plates that contained faulty pDNA were reordered/remade and subjected to the same QC process detailed above. pDNA preparations with the correct sequence but contaminations greater than 1% of another CFTR variant plasmid were recloned, re-amplified and subjected to the described rigorous QC.

### Cell Culture

The original unmodified FRT cell line, which does not express endogenous CFTR, was grown and maintained in culture flasks containing Ham’s F-12 nutrient mixture (Coon’s modification, F6636, Sigma-Aldrich, USA) supplemented with 10% FBS (SH30066.03, Hyclone, USA), 1% Penicillin-Streptomycin (15070063, Gibco^TM^, USA), and sodium bicarbonate (S8761, Sigma-Aldrich, USA) at 37°C in a 5% CO_2_/95% air atmosphere.

For pDNA transfection, FRT cells were trypsinized (0.25% Trypsin EDTA, 25200056, Gibco^TM^, USA) at about 80% confluency and transiently transfected using a reverse transfection protocol. Per transfection condition, 250 µl of 3.5 x 10^5^ cells/ml were added to a 1.5 ml Eppendorf tube containing a premixed solution of 50 μl OptiMEM media (31985-088, Gibco^TM^, USA), 0.4 μg of CFTR wild-type or variant pDNA, 0.75 μl Lipofectamine LTX and 0.6 μl Plus reagent (15-338-030, Thermo Fisher Scientific, USA). After 30 minutes incubation, an aliquot of the transfection mix containing about 87,500 cells was transferred onto each filter of a Costar 24-well Transwell® high-throughput screening (HTS) filter plate (3378, Corning, USA) according to the plate layout shown in **Figure 1A**. 750 µl of F-12 Coon’s media (10% FBS) was added to the bottom well of the filter plate. Two days post seeding, 150 μl of top and 350 μl of bottom media was replaced with fresh F-12 Coon’s media (10% FBS). On day 4 (24h prior to assay), top and bottom media were replaced with 250 μl and 750 μl of F-12 Coon’s (10% FBS), respectively, containing either vehicle (DMSO) or a combination of CFTR correctors: 3.5 μM tezacaftor (TEZ, VX-661, AA Pharma, USA) and 3 μM elexacaftor (ELX, VX-445, Dipharma, formerly Kalexsyn, USA).

**Figure 1:**
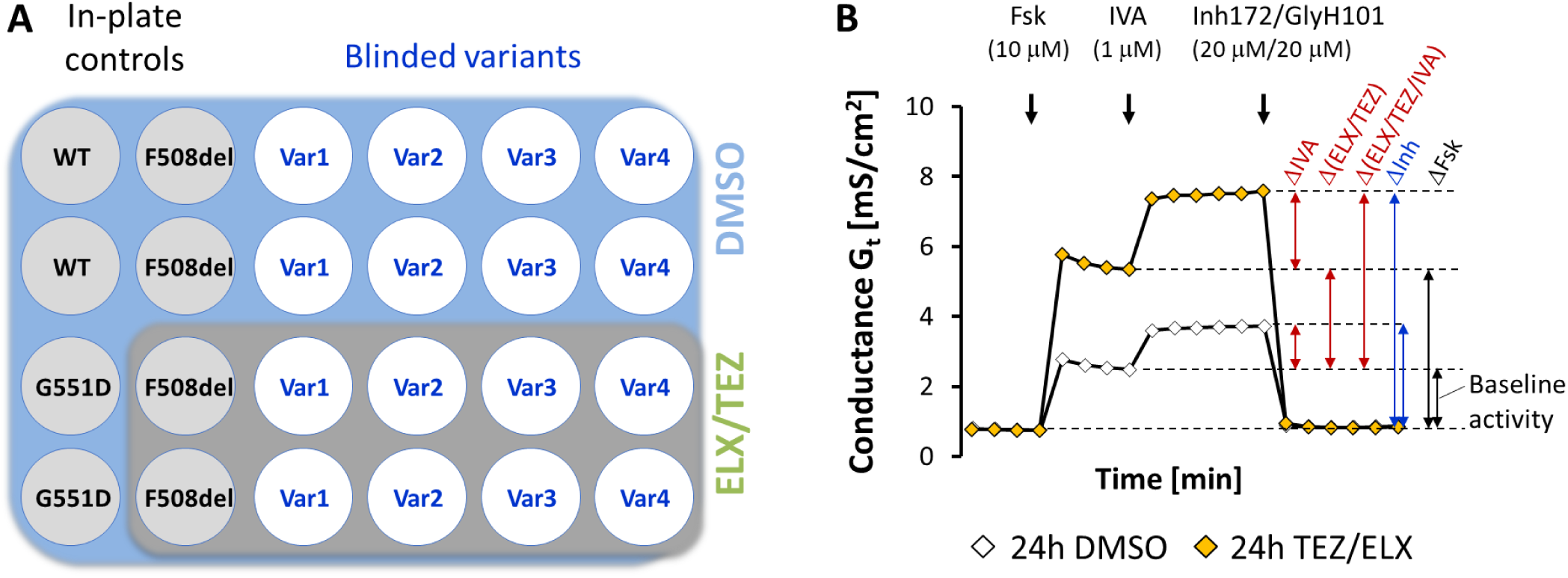
(**A**) Plate layout of a Costar 24-well Transwell® high-throughput screening (HTS) filter plate used in the FRT cell transepithelial conductance assay. Wells inside the dark grey background were treated with CFTR modulators ELX/TEZ. All other filter cultures were treated with vehicle (DMSO) as described above. (**B**) Representation of two separate conductance traces at 5 min. sampling rate of a corrector (ELX/TEZ) and potentiator (IVA) responsive CFTR variant as observed in two different wells: one treated for 24h with DMSO (white diamonds) and one with ELX/TEZ (orange diamonds). After recording the resting conductance, forskolin (Fsk) is added to each well to stimulate CFTR activity followed by addition of potentiator IVA to test the variant response to IVA in the presence and absence of correctors. Addition of CFTR specific inhibitor cocktail completes the assay. Variant responses to modulators IVA, ELX/TEZ, and the triple combination ELX/TEZ/IVA are indicated by the red arrows (Gt change from Fsk baseline).

### Electrophysiology

For data QC and normalization of CFTR variant function, WT CFTR and CFTR bearing the c.1652G>A (p.Gly551Asp; legacy: G551D) or c.1521_1523del (p.Phe508del; legacy: F508del or [delta]F508) variants were included on every 24-well Transwell® HTS assay plate. Four CFTR variants were evaluated on each plate as shown in **Figure 1A**.

One hour before the functional assessment of CFTR variants, cells were switched from growth media (Coon’s F-12, 10% FBS) to 20 mM HEPES buffered (pH 7.4) Coon’s F-12 media (without FBS and bicarbonate) and incubated on the assay platform in temperature-controlled plate holders or in a CO_2_ free incubator at 37°C. Transepithelial conductance (G_t_) measurements were made using an automated 24-channel current clamp system (TECC-24; EP Design, Belgium). Current responses to a sine wave at 1 Hz and an amplitude of 10 mV were taken at regular intervals, i.e. about every 5 minutes at CF and RF and 0.5 minutes at UT to measure transepithelial conductance G_t_. Pre-forskolin resting conductance was measured for 15 minutes, then cells were stimulated with 10 μM forskolin (F6886, Sigma-Aldrich, USA) followed by 1 μM ivacaftor (IVA, VX-770, AA Pharma, USA). Lastly, CFTR was inhibited with a combination of 20 μM CFTR_Inh_-172 (016665, Matrix Scientific, USA) and 20 μM GlyH101 (5485, Tocris Bioscience, USA). Reagents were added at 20 ± 5 minutes intervals and the change in conductance (ΔG_t_) was calculated for every reagent addition (see **Figure 1B**).

### Western blots

For assessment of CFTR protein maturation, transiently expressing FRT cells were harvested post electrophysiological assay after addition of Pierce IP Lysis Buffer (87788, Thermo Fisher Scientific, USA) or freshly made lysis buffer (50 mM Tris pH 7.4, 150 mM NaCl, 1 mM EGTA, 1 mM EDTA, 1% Trition X-100), Roche cOmpleteTM, EDTA-free protease inhibitor cocktail (4693159001, Sigma-Aldrich, USA), and rocking at 4°C for 30-60 minutes.

UT: The insoluble fraction of the lysate was removed via centrifugation through 1 mm glass fiber Acroprep Advance 96-well filter plates (8031, Pall, USA). The soluble flow-through fraction was analyzed by SDS-PAGE. Core (high mannose)-glycosylated CFTR (Band B), consistent with localization to the endoplasmic reticulum, and complex glycosylated CFTR (Band C), expected for functional, plasma membrane-located protein, was visualized by adding CFTR monoclonal antibody 596 (1:10000, J. Riordan and M. Gentzsch [7], University of North Carolina, Chapel Hill, NC) and anti-mouse IgG secondary antibody IRDye 800CW (1:15000, 926-32210, LI-COR, USA) and quantified using the LI-COR (Lincoln, NE) Odyssey CLx Imaging system and software Image Studio Version 5.

CF: The insoluble fraction of the lysate was pelleted via centrifugation for 20 minutes at 21,100 x g (4oC). The supernatant was analyzed by SDS-PAGE and Western blot or the WES capillary electrophoresis system (ProteinSimple, USA. For SDS-PAGE, CFTR was visualized using as above the antibody 596 at 1:2000 and goat-anti-mouse IgG HRP secondary antibody (1:10000, 115-035-166, Jackson Immuno Research Labs) and quantified using the volume tool in Bio-Rad Laboratories Image Lab 4.0 software. For WES, CFTR was visualized with the primary monoclonal antibody 450 (1:4000, J. Riordan and M. Gentzsch [7], University of North Carolina, Chapel Hill, NC) and ProteinSimple anti-mouse detection module (DM-002), using the 66-440 kDa separation module with 8 X 25 capillary cartridges (SM-W008, ProteinSimple, USA). The CFTR signal was quantified with the ProteinSimple analysis software Compass for SW 4.0.0. CFTR variants (Q1209P, M1210K) with mutations in the epitope of the 596 antibody were detected with the 450 antibody.

### Data processing and analysis

#### Filtering of functional data

Prior to averaging the data, the following exclusion criteria were applied to meet internal quality standards:

1. Data from Transwell® filters with transepithelial electrical resistance (TEER) values below 200 Ωcm^2^ were excluded from the analysis.
2. Data from entire 24-well assay plates were excluded from analysis, if more than 1/3 of wells had TEER values below 200 Ωcm^2^ (total of nine assay plates).
3. Conductance traces with only partial response to addition of CFTR inhibitor cocktail Inh-172/GlyH101 were removed from the analysis.
4. Three assay plates at one of the sites had both WT controls failing the QC criteria outlined above, so %WT metrics could not be computed, and thus were excluded from the analysis.
5. Upon initial measurement a few electrophysiological traces were considered of questionable quality and were repeated (at the same site); in cases where duplicate measurements were available *and* clear difference in trace quality was observed, only the higher quality data was included for analysis (regardless of the consistency between the repeats or with the other sites).

In summary, a total of 13,714 wells were measured at the three participating sites. Six-hundred and forty-five wells were removed based on the specific and formal criteria 1-5 listed above. Independently, 348 wells from replicate measurements available at individual sites were manually flagged as poor quality; 194 of those (56%) also did not pass the formal QC criteria 1-5, while the remaining 154 wells were manually removed due to various other issues. Thus, 645+154=799 individual wells (5.8% fail rate) were removed from the data across all three participating sites.

### No filtering was performed for the Western blot data

#### Normalization of functional data

The raw conductance (G_t_) time course data recorded from each well by TECC instruments in all participating laboratories were centrally processed using custom software developed in-house in R language (Ref [8]) (source code available upon request). The G_t_ responses to acute addition of each compound (forskolin (ΔFsk), ivacaftor (ΔIVA), inhibitor (ΔInh)) were calculated as the difference between G_t_ on the respective established plateaus as illustrated in **Figure 1B**. The measured response values ΔFsk and ΔFsk+ΔIVA from each well of a 24-well assay plate were further normalized to the WT function calculated as average ΔFsk value from the two vehicle-treated WT intraplate control wells (**Figure 1A**), giving Fsk %WT and Fsk+IVA %WT metrics.

#### Normalization of Western blot data

Protein samples from each well of a 24-well assay plate were run on the same gel. Background-subtracted values for CFTR band B and band C intensities were used to (1) calculate average band C intensity, total protein (band C + band B), and band C/band B ratio in intraplate WT control wells; (2) normalize the metrics from all wells on the plate to the respective WT average, resulting in band C %WT, C/B %WT, Total protein %WT; additionally, band C %Total was computed for each well (using the total for that same well).

#### Outlier detection and validation

To quantify the consistency of the electrophysiology measurements across the laboratories for each individual pDNA clone and treatment condition (vehicle or modulators), the measurements at the three different sites were compared by applying a moderated ANOVA test followed by post-hoc Tukey pairwise HSD calculation (see **Supplement** for details). The analysis was applied separately to each of FSK %WT and FSK+IVA %WT metrics, in DMSO or with ELX/TEZ (over 2600 measurements total), and indicated that in ∼70% of all cases the metrics were consistent across all three labs, in 25% of cases one site generated an inconsistent metric (while the other two were consistent), and in the remaining 5% of cases a metric was inconsistent across all three reporting labs according to the chosen criteria.

Based on the outlier analysis, additional validation has been performed by choosing 95 pDNA clones showing responses measured at CF, or RF, or both that were among the least consistent with the measurements performed at the other site(s) (smallest p-values). Of those variants, 55 were assayed again at CF, 31 at RF, and 9 both at CF and RF. The putative outliers exhibited a trend for improvement but were not fully resolved after the repeated measurement (see **Supplement**), thus there were no objective and statistically sound criteria to discard the outlier observations in the primary screen and they were included. In the following analysis the primary and validation data are simply combined (resulting in larger numbers of assay wells/assay plates for some variants) and averaged as described below.

#### Calculating average variant response

After normalizing the per-well measured values as described above, all the metrics were averaged per each CFTR variant and treatment across in-plate replicates, repeated plates, different pDNA clones (in a few cases where multiple, independently generated clones were available for the same variant), and across the three sites. Since the variances of the measured values due to those multiple factors are intrinsically different, we applied a mixed effect model to each variant, treating repeated plates and different sites as random effects (see Supplement for details).

#### Empirical Bayes method

Variant responses (%WT) to ELX/TEZ/IVA were averaged over all three labs and the averaged responses served to calculate a standard error (SE) estimate for each variant. Using this collection of sample average responses and SEs, empirical bayes was used to fit a posterior distribution for the true response. A Gaussian prior was used with the mean and variance estimated from the data. Sample average responses were assumed to have gaussian likelihoods, with variance heteroskedasticity allowed. Models were fit using the package “ebnm” (version 1.0.9) in R. This approach allows to calculate both posterior point probabilities that the true underlying response for any given variant exceeds a specific cutoff and their credible intervals. This facilitates an evaluation of uncertainty, though there are relatively strong modeling assumptions.

## Results

### Pharmacological validation of the transiently expressing FRT cell model

The goal of this study was to evaluate whether ELX/TEZ/IVA increased Cl^-^ transport for a large number of CFTR variants. In the absence of tissue samples and to avoid the time required for production of hundreds of stable cell lines, an alternative approach was developed. FRT cells were reverse transfected with variant plasmid and seeded onto permeable supports. Four days later, cells were treated with either DMSO (vehicle) or a combination of TEZ and ELX. After 24 hours of incubation, the resting (pre-forskolin) transepithelial conductance (G_t_) and G_t_ changes in response to the addition of forskolin, IVA and a cocktail of inhibitors were measured as outlined in **Figure 1B**. This validation section focuses on WT CFTR and the variants F508del and G551D, both of which have well established responses to CFTR modulators (Van Goor+, 2009 [9], Van Goor+, 2011 [10]) and were used as controls on all assay plates. Each mean value and standard deviation in this section is based on over 1100 individual measurements conducted throughout the study. Responses to inhibitor, ΔG_t_ Inh, are given as absolute values.

For cells transfected with WT CFTR, the resting G_t_ was 0.8 ± 0.6 mS/cm^2^. The addition of forskolin increased the G_t_ by 10.7 ± 3.8 mS/cm^2^, a ∼13-fold increase. IVA further increased the forskolin-stimulated G_t_ by 1.8 mS/cm^2^. The addition of inhibitors fully reversed the increase in G_t_ caused by forskolin and IVA, decreasing G_t_ by 12.3 ± 4.4 mS/cm^2^ to the original baseline conductance (**Figure 2A**).

**Figure 2:**
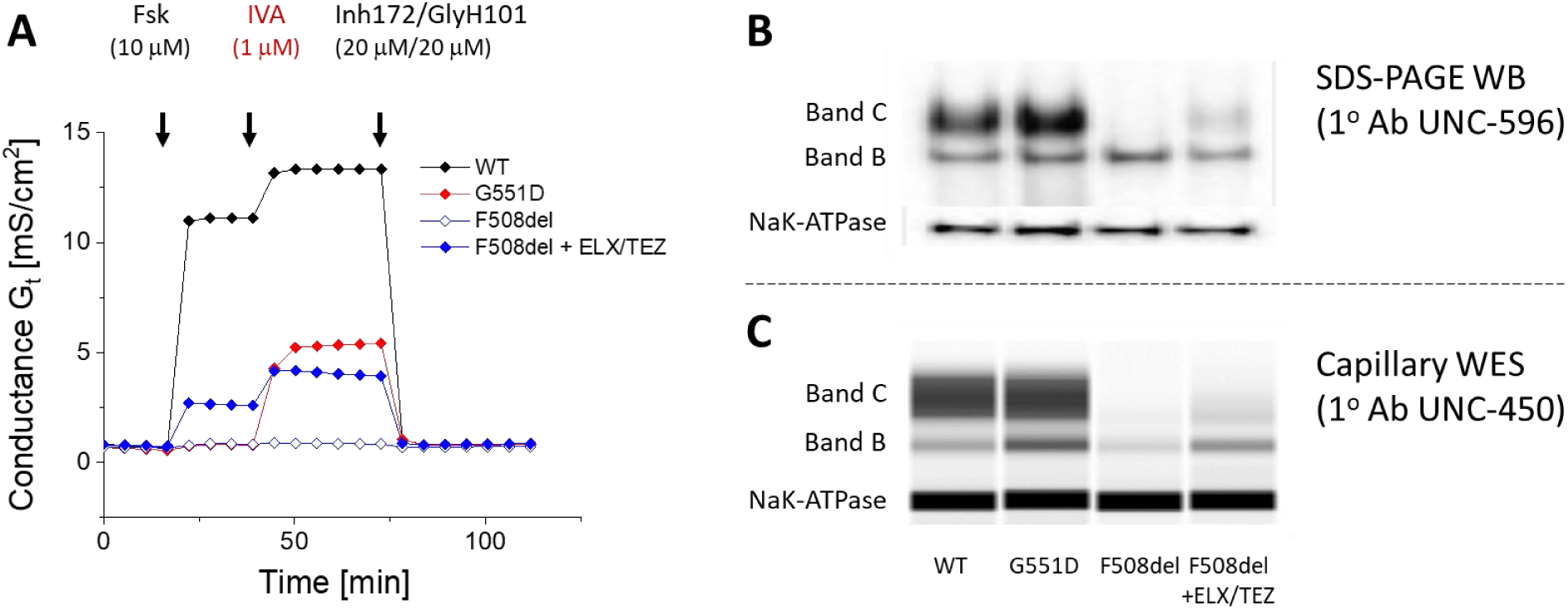
(**A**) Representative recordings of transepithelial conductance (G_t_) and G_t_ responses to the addition of forskolin, IVA and a combination of inhibitors from electrically tight (>200 Ωcm^2^) FRT cell monolayers grown on Transwell® permeable filter supports 5 days after reverse transfection of cells with WT, G551D, and F508del CFTR pDNA. Cells were treated for 24h with DMSO (WT, G551D, F508del) or ELX/TEZ (F508del). Representative images of Band B and mature Band C CFTR as determined by SDS-PAGE (**B**) or capillary Western blot (**C**) techniques for corresponding CFTR variants and treatments.

The resting G_t_ in cells transfected with F508del CFTR was 0.7 ± 0.6 mS/cm^2^. The ΔG_t_ response to the addition of forskolin and IVA to untreated F508del CFTR cells was 0.2 ± 0.2 mS/cm^2^ and 0.0 ± 0.1 mS/cm^2^, respectively. The resting G_t_ in F508del CFTR cells treated overnight with ELX and TEZ was 0.8 ± 0.6 mS/cm^2^. Stimulation with forskolin increased G_t_ by 1.6 ± 0.8 mS/cm^2^ and G_t_ was further increased by 1.3 ± 1.1 mS/cm^2^ by the addition of IVA. The forskolin plus IVA G_t_ values were close to 30% of the G_t_ of forskolin-stimulated WT CFTR transfected cells. Increases in G_t_ were fully reversed through the addition of the inhibitor cocktail (ΔG_t_ Inh = 2.9 ± 1.4 mS/cm^2^, **Figure 2A**).

The resting G_t_ values in cells transfected with G551D CFTR was 0.7 ± 0.7 mS/cm^2^. Forskolin caused only a small increase in G_t_ (0.7 ± 0.9 mS/cm^2^). In contrast, IVA caused a ∼5-fold further increase in the forskolin-stimulated G551D CFTR cells (3.6 ± 2.1 mS/cm^2^). The forskolin plus IVA G_t_ values were approximately 40% of the G_t_ of forskolin-stimulated WT CFTR transfected cells. The increases in G_t_ were completely reversed by the inhibitor cocktail (ΔG_t_ Inh = 4.1 ± 2.7 mS/cm^2^, **Figure 2A**).

The above phenotypes and pharmacological profiles are the expected results for cells expressing WT, F508del CFTR and G551D CFTR. Western blots (**Figure 2B** and **2C**) from lysates of the above cells documented the expression of CFTR. In the case of WT CFTR and G551D CFTR transfected cells, Band C was the dominant band observed. Band B was the dominant band in untreated F508del CFTR cells. Treatment of the F508del CFTR cells with ELX and TEZ caused an increase of Band C relative to Band B, consistent with the phenotypical and pharmacological results above. With this confirmation and reproduction of established functional and biochemical responses, the assay is considered validated and suitable for theratyping.

### Correlation of CFTR variant functional responses between sites

CFTR WT and 655 variants (608 at UT, 654 at RF and 655 at CF) were screened in the FRT cell line to assess variant baseline activity and variant responsiveness to IVA alone and ELX/TEZ/IVA triple treatment. **Figure S1** shows robust correlations (Pearson correlation coefficients of 0.8 – 0.9) for CFTR variant baseline activities and variant responses to addition of ELX/TEZ/IVA between sites.

**Figure 3** provides a different view of the data congruence among the three sites. For the baseline conductances and for the responses to triple treatment measured at each lab, the linked bar charts or alluvial plots, show the agreement between the sites when binning the conductance responses into three bins, <10%, 10% – 50%, and >50% of wild type function. The white stacked bar in the middle of each graph shows the distribution for CF and the bars on the left and right represent the distributions at UT and RF, respectively. The colored connections in between illustrate the pairwise data correspondence between CF and the other two labs, with the connector colors corresponding to CF measurements.

**Figure 3:**
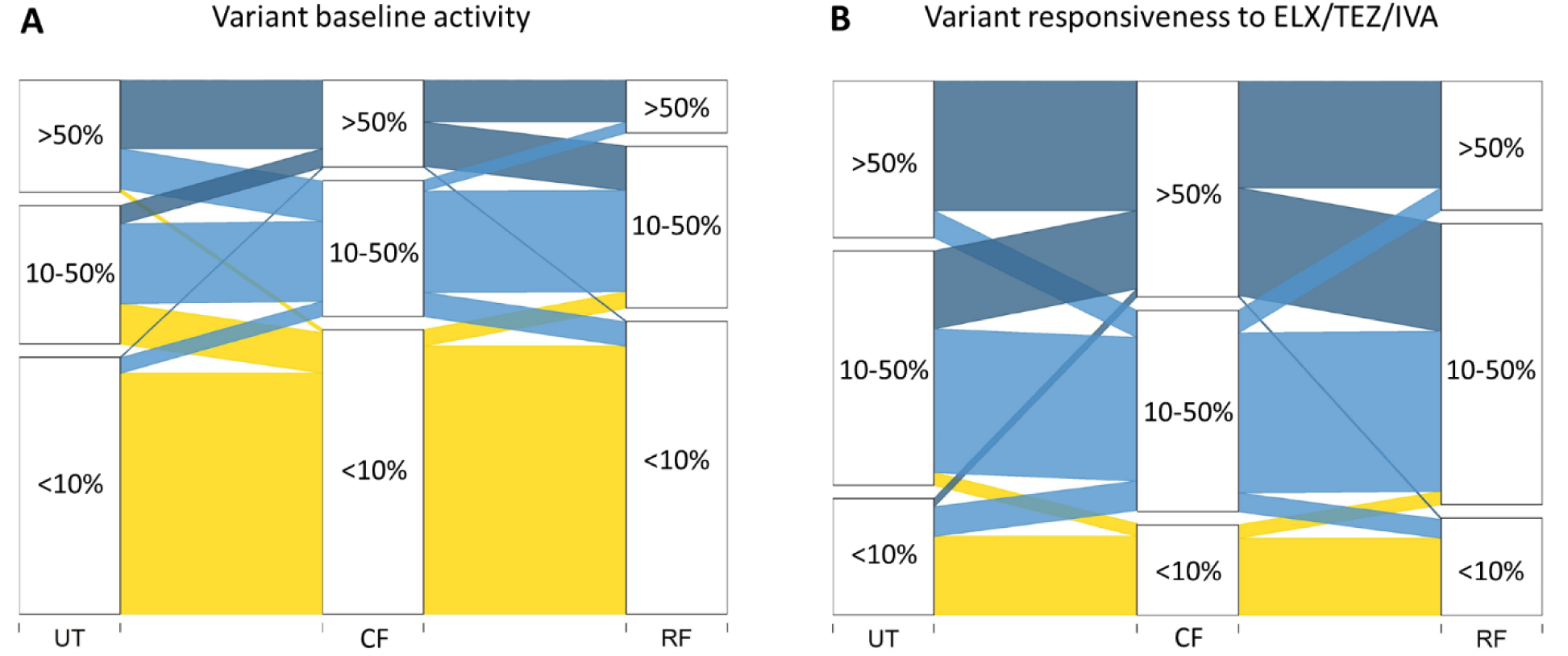
Linked bar charts illustrating the data congruence between CF and RF, UT sites with respect to CFTR variant baseline activity (**A**) and responsiveness to ELX/TEX/IVA (**B**). Connecting bars are colored based on CF site measurements (yellow: < 10% of WT function; light blue: 10%-50%; dark blue: >50%)

A quantitative summary of the congruence of measurements among the three sites is provided in **Tables 1A** and **1B**. After excluding F508del, G551 and WT controls, measurements for 606 (baseline function) and 605 (functional response to ELX/TEZ/IVA) of the 653 remaining CFTR variants were considered because data exists from all three labs. Agreement among responses is summarized in **Table 1** using the binary classification of ‘above’ or ‘below’ the 10% WT threshold. For example: of the total 109 variants, unapproved and approved, that based on CF measurements improved <10% relative to WT function upon ELX/TEZ/IVA treatment, 86 (79%) also had <10% increase relative to WT function at the other two labs, and only 7 variants (6%) were measured with ≥10% improvement relative to WT function at both other labs. Of the 496 variants with ≥10% of WT function improvement after ELX/TEZ/IVA treatment measured at CF, 438 (88%) also showed ≥10% improvement relative to WT function at both other labs, and only 11 (2%) had <10% WT function improvement at both other labs.

**Table 1:**
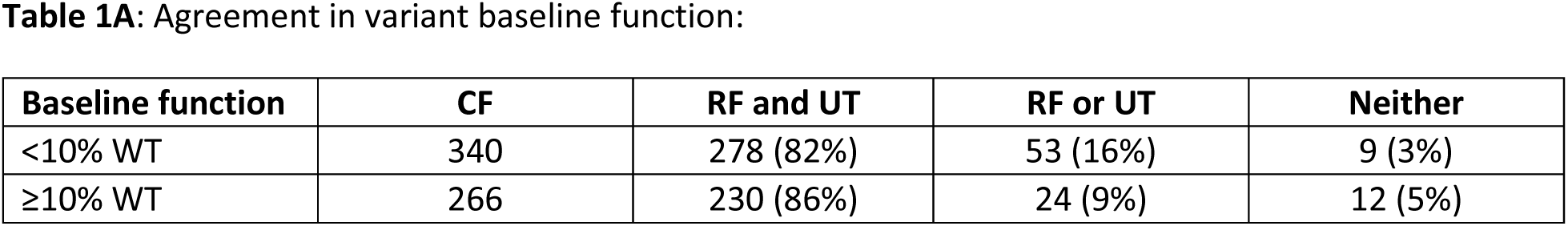
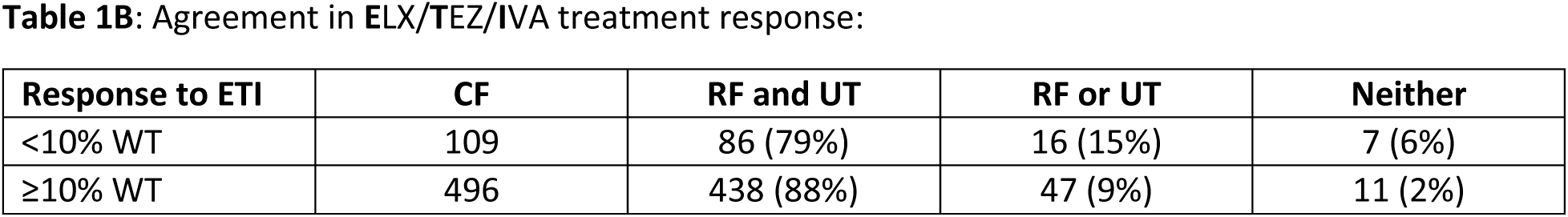
Number of labs that agree with CF classification (column 1) for CFTR variant baseline function (**1A**) or CFTR variant responsiveness to ELX/TEX/IVA (**1B**). Note percentages are computed out of samples that had a valid response at all 3 labs, i.e., 47 of 653 variants were left out of the CFTR variant baseline comparison, and 48 were left out of the comparison of CFTR variant response to ELX/TEZ/IVA.

### Most variants respond to ELX/TEZ/IVA triple modulator combo with an increase in CFTR chloride conductance

For a representative variant subset, **Figure 4A** shows variant baseline activity, i.e., forskolin-only response, which is associated with disease liability, and variant responsiveness to forskolin plus ELX/TEZ/IVA, which is expected to correlate to a potential clinical improvement. Also included are the individual variant responses to forskolin plus IVA and forskolin plus ELX/TEZ (for the complete dataset, see **Table S1**). All data are normalized to the intraplate control WT forskolin response (100%) as described in the methods section. CFTR variants are sorted by their functional response to forskolin. Benchmark levels for G551D + IVA and for F508del + ELX/TEZ/IVA are also indicated. In this assay, a lumacaftor/ivacaftor treatment of F508del resulted in ∼10% of WT function (data not shown).

**Figure 4:**
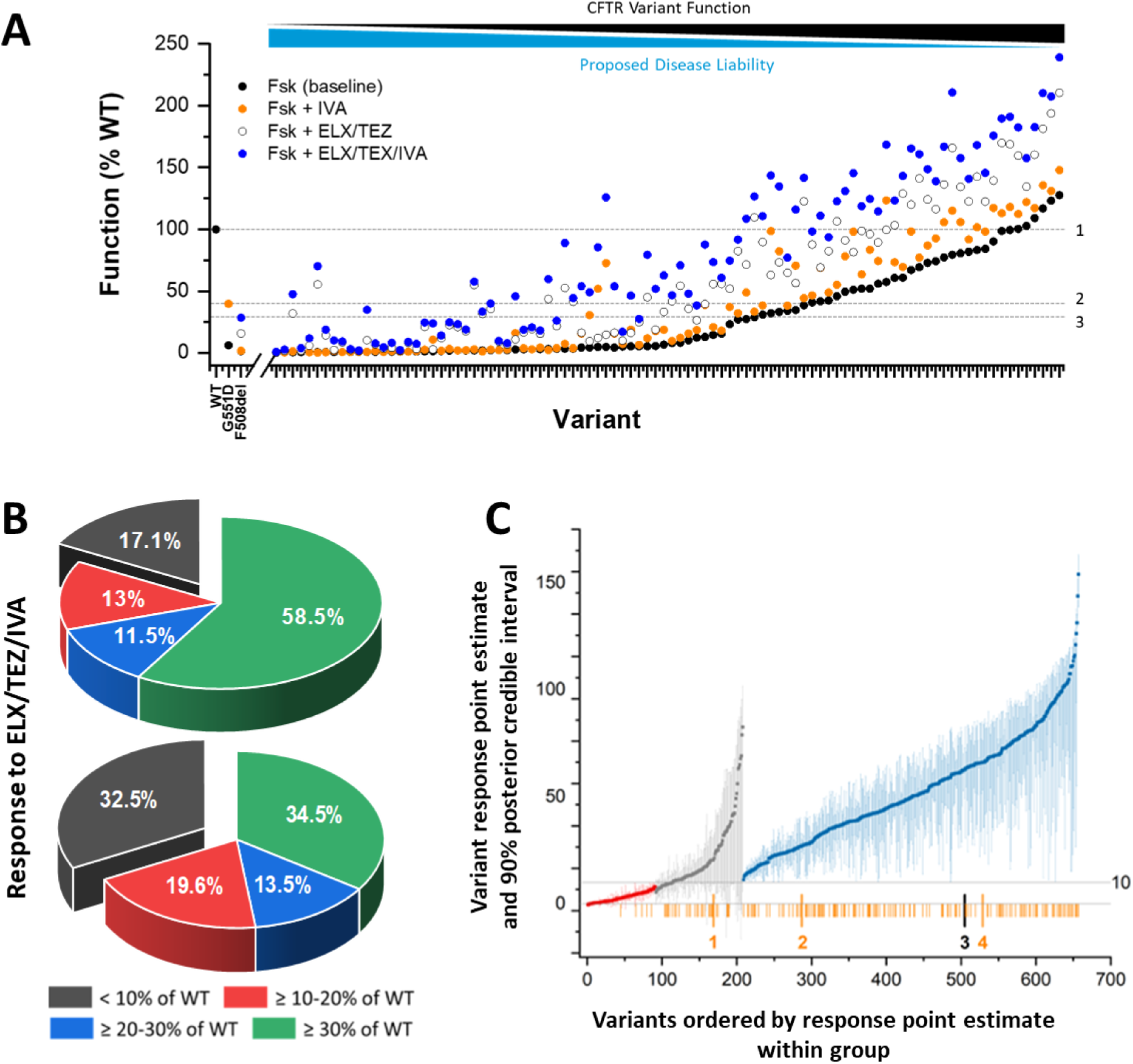
(**A**) Example of a representative subset of CFTR variants and their respective responses to forskolin (Fsk, black dots), Fsk plus CFTR potentiator IVA (orange dots), Fsk plus CFTR modulators ELX/TEZ (open circles), and Fsk plus the triple combination ELX/TEZ/IVA (blue dots). Dashed lines indicate benchmark values for: 1 = WT (100%, n = 1122), 2 = G551D + IVA (39.8%, n = 1132), 3 = F508del + ELX/TEZ/IVA (28.5%, n = 1124). (B) Responsiveness of CF-associated variants to ELX/TEZ/IVA: color-coded ranges represent improvement in CFTR function with ELX/TEZ/IVA over baseline (ΔFsk). **Top**: Percentage regardless of variant baseline function. 82.9% of 655 variants show an increase in CFTR function of ≥10% of WT. **Bottom**: Percentage for subset of 342 variants with baseline function <10% of WT CFTR. 67.6% of all variants with <10% of normal baseline function show an increase in CFTR function of ≥10% of WT. (C) Graph showing point estimates and 90% posterior credible intervals for the true underlying response for 655 CF-associated variants to ELX/TEZ/IVA. Variants are ordered by their response point estimate. The variants in blue have their lower bound ≥10, those in red have their upper bound <10, and those in grey have their upper/lower bound cross the 10% WT threshold. Vertical lines in orange indicate Trikafta® approved variants on the x-axis. Lines with numbers highlight x-axis positions of included controls: (1) F508del (blinded samples), (2) F508del, (3) WT, and (4) G551D in-plate controls.

The majority of the 655 known or putatively CF-associated variants included in this study have little to modest baseline activity and are likely disease-causing variants. For example, 342 variants display an averaged baseline activity of less than 10% of WT function in our assay and 474 less than 30% function relative to WT. One hundred and four variants in this study had baseline activity of >50% WT in the FRT cell assay, which may suggest that they are not disease causing by themselves (in the context of the DNA sequence utilized).

Overall, the results from this *in vitro* study suggest many rare variants in the evaluated set are disease causing and correctable by modulator treatment. Nearly 83% of all CFTR variants tested responded to ELX/TEZ/IVA triple modulator treatment with an increase in chloride ion transport activity of 10% of WT function or more (**Figure 4B**, upper pie chart: sum of red, blue, and green percentages). Based on data from a similar assay with stable FRT cell lines, variants that improved by at least 10 percentage points of WT function in response to ELX/TEZ/IVA treatment have been classified as responders and were approved for Trikafta® therapy. By applying the same threshold to the data from this study, there is excellent agreement with the FDA label for Trikafta®. Ninety-four percent of approved variants were also found to be responsive in the study reported here.

**Figure 4B**, lower pie chart, shows a graphical representation (percent of total) of the mean variant response to ELX/TEZ/IVA for the 342 variants with less than 10% normal (baseline) function associated with a higher disease liability. Based on the determined values for responsiveness, 67.6% of these variants responded to treatment with ELX/TEZ/IVA with an increase in CFTR chloride conductance of ≥10% of normal and 34.5% have a mean response of ≥30% of normal. 32.5% of variants responded with less than 10% WT functional improvement when treated with the triple modulator combo.

This rather binary view of the data, however, neglects any uncertainties associated with the determined individual (mean) variant responses to ELX/TEZ/IVA. In a separate analysis, an empirical bayes method was used to evaluate the likelihood for each of the 655 known or putatively CF-associated variants included in this study that the true underlying response is ≥10% of WT function (Figure 4C).

Based on the above described empirical bayes method, about 68% of CF-associated variants (Figure 4C, blue dots) classify as responders to treatment with the triple modulator combo ELX/TEZ/IVA given a lower bound of ≥10% of WT and a 90% credible interval. Though there is a clear cut-off (red vs grey vs blue colored variants), variants with a modulator response close to the 10% WT threshold (e.g., N1303K = 9.4%), could be classified as responders or non-responders, depending on the analytical method applied. This holds especially true for statistically low-powered studies. Here, the 655 CF-associated variants were binned into 3 categories (responder, non-responder, and borderline responsive) based on their 90% posterior credible intervals for the true underlying *in vitro* response to ELX/TEX/IVA (Table 2).

**Table 2:**
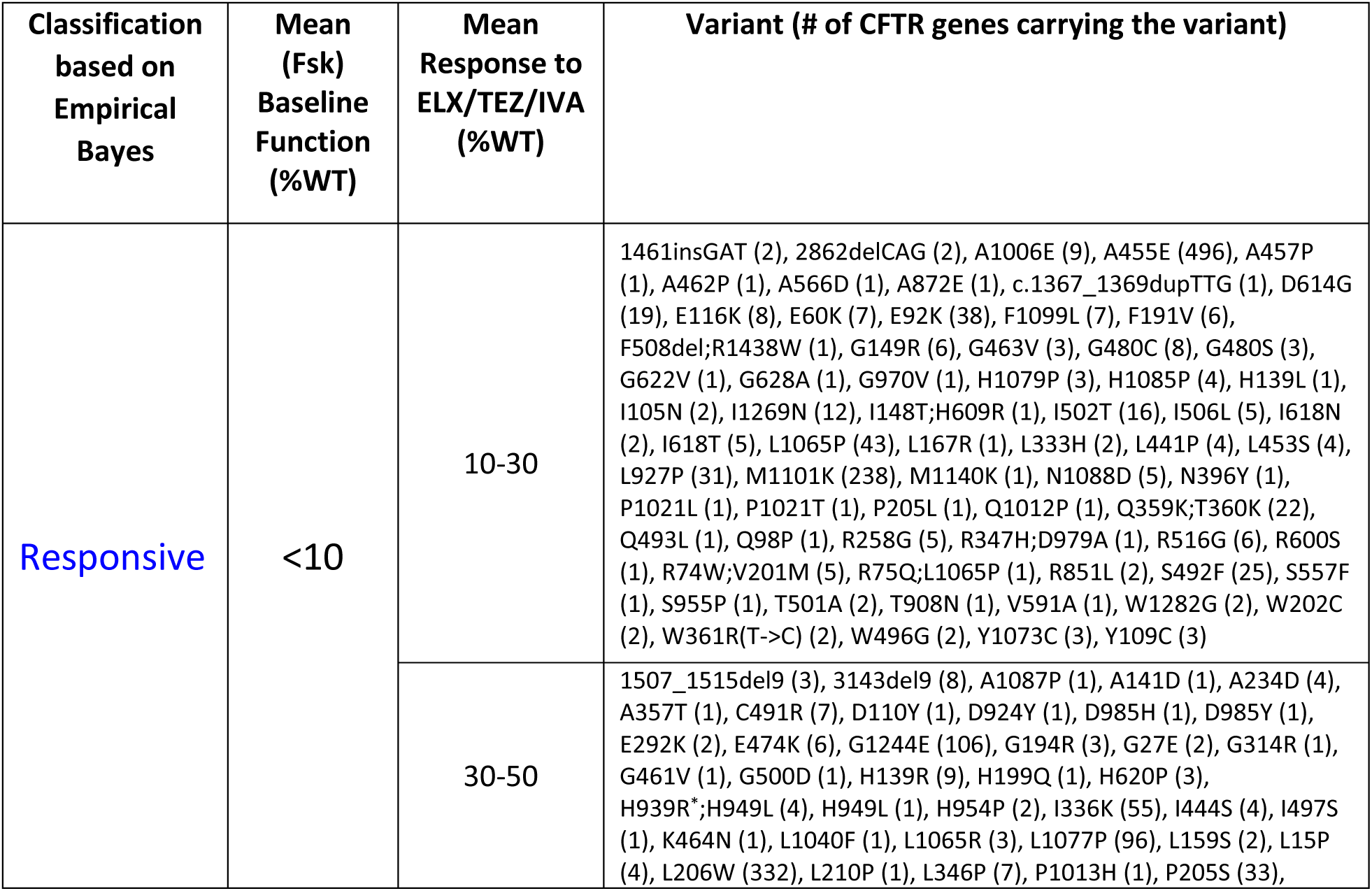
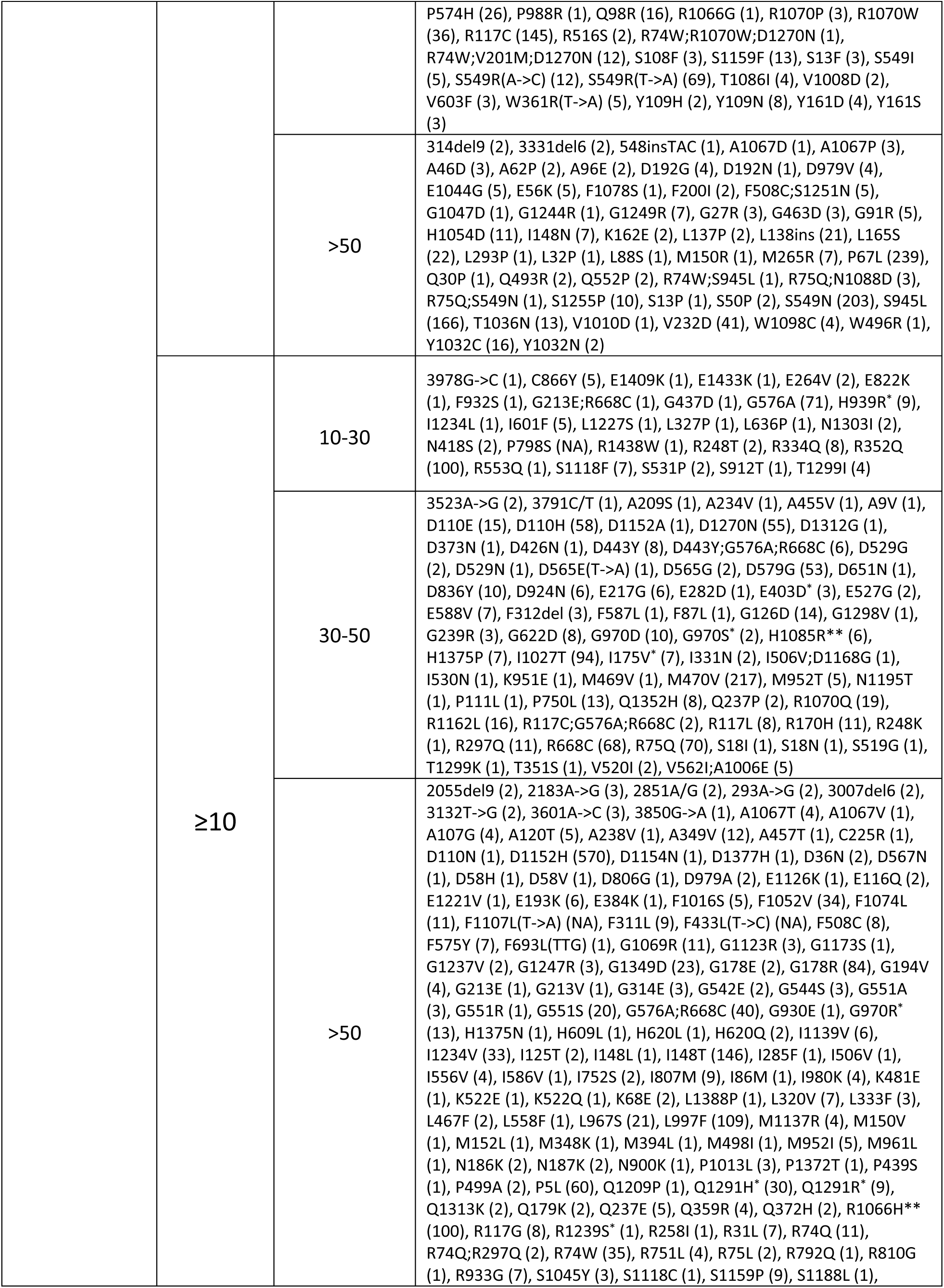
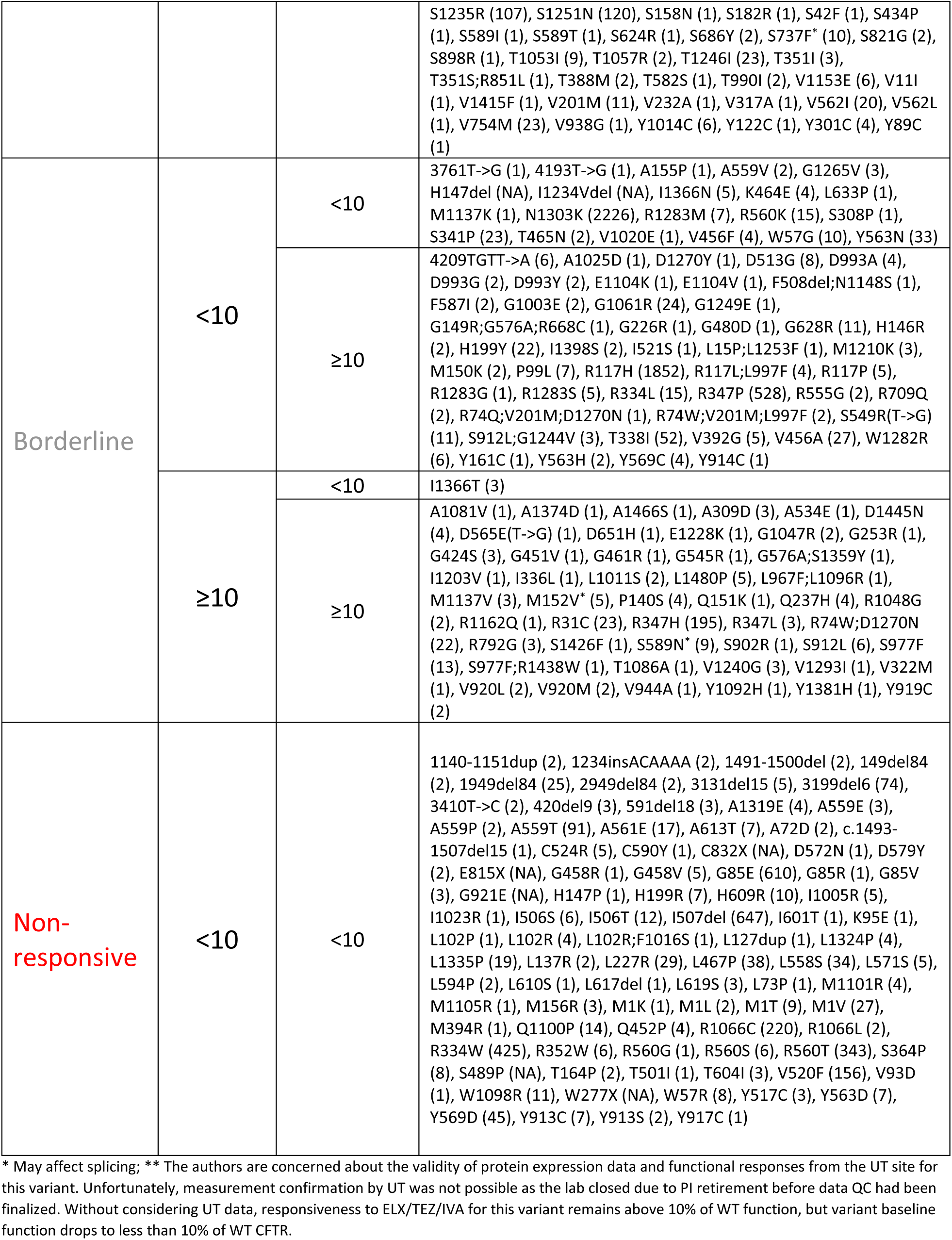
Variant Response Size to ELX/TEZ/IVA Treatment in %WT Function

### Many CFTR variants not yet FDA approved are responsive to ELX/TEZ/IVA combination

The functional analysis of 655 CF-associated CFTR variants in the FRT cell model used in this study yielded 231 CFTR variants (Figure 5, triangles in green box) with baseline CFTR function of <10% of WT that respond to ELX/TEZ/IVA by an increase in CFTR function ≥10% of WT. 152 of those CFTR variants (red triangles in green box) are not currently approved for the highly effective modulator therapy Trikafta® (for a complete list of the 178 Trikafta®-approved variants (cf. FDA, 2021rev [3]).

**Figure 5:**
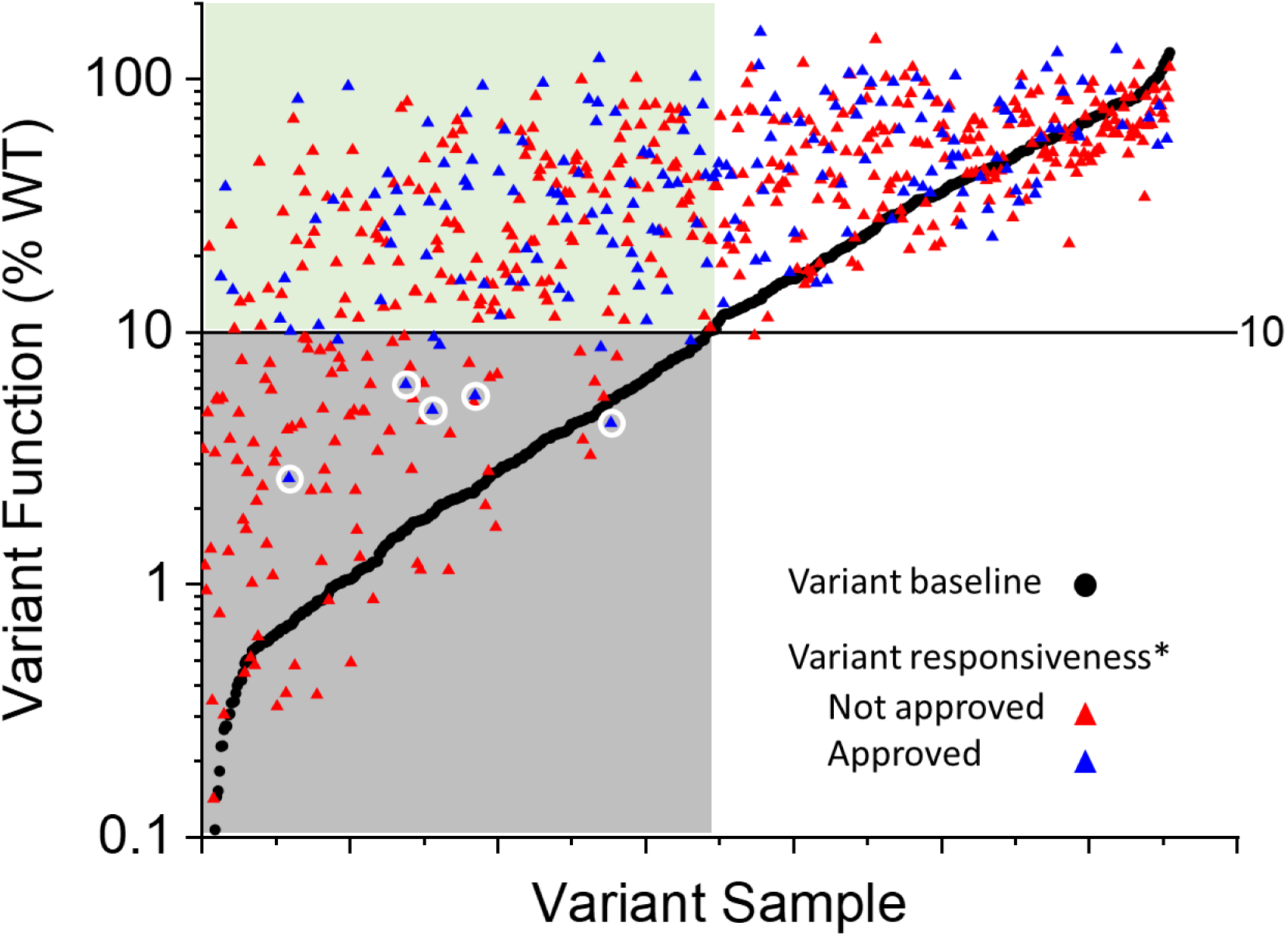
Baseline activity and responsiveness to ELX/TEZ/IVA for all 655 known and putatively CF-associated variants are plotted. Each black dot represents the mean baseline function of a specific CFTR variant and each of the triangles represents the corresponding responsiveness of a variant to ELX/TEZ/IVA. Black dots in the grey box represent variants w/ baseline activities of less than 10% of WT. Triangles in the green box represent variants that respond to combined modulator treatment by an increase of 10% of normal or more. Blue triangles represent the 174 (out of 178) Trikafta approved variants screened in our study. Circled are five variants that are approved for Trikafta® therapy but clearly missed the responsiveness threshold in this study. **\***Responsiveness = change from Fsk (DMSO) baseline, i.e., variant function (%WT) in the presence of ELX/TEZ/IVA - variant (Fsk) baseline function (%WT).

We also identified variants with borderline responsiveness to ELX/TEZ/IVA, i.e. those that missed or made the arguable cut-off of ≥10% improvement in CFTR function by a small margin. Among those that fell short (by less than 1.5% of WT function), solely judged by their mean responsiveness, are five variants already approved for modulator therapy: S341P (9.1%), V456F (9.4%), Y563N (9.3%), R1283M (8.8%), I1366N (8.7%). Only five of the 174 Trikafta approved variants tested in this study clearly missed the ≥10% response threshold in our assay (**Figure 5**, circled triangles): G85E (5.6%), R352W (4.3%), S364P (4.9%), L1324P (2.6%), L1335P (6.5%). However, at higher (10 μM) ELX/TEZ concentration, compared to standard 3 μM ELX and 3.5 μM TEZ used throughout the study, the responsiveness for CFTR variants L1324P, V456F, and R1283M improved from 2.6% to 10.4%, 9.3 to 22.2%, and 8.8% to 29.2%, respectively (data not shown). No improvement was observed at higher ELX/TEZ concentrations for the other seven variants mentioned above. In addition, despite acute in-assay additions of 1 μM IVA, which is 10 to 40-fold higher than the reported EC_50_ values for IVA in FRT cells expressing G551D or F508del, respectively (Van Goor F+, 2009 [9]), the maximum potentiator response may not have been achieved for variants with an extreme right-shifted IVA concentration-response characteristic (± ELX/TEZ).

## Summary and Discussion

In total, 655 known or putatively CF-causing CFTR variants were assessed for their responsiveness to the triple drug combination ELX/TEZ/IVA, 478 of which are not currently approved for any CFTR modulator therapy. Three hundred seventy-six of these 478 variants had increased Cl^-^ transport function by ≥10% of WT after treatment with ELX/TEZ/IVA, regardless of their baseline function.

The initial *in vitro* studies supporting the inclusion of responsive variants on the FDA drug label focused on the relatively more common CFTR variants. The goal of this study was to assess the ELX/TEZ/IVA modulator responsiveness of rare variants in the CFTR2 database suitable for a cDNA model, even if reported only in a single pwCF. Notably, 152 additional CFTR variants were identified that had less than 10% of WT function at baseline (generally associated with disease) that improved by more than 10 percentage points of wild type function in response to drug treatment. This level of response is an indicator of probable clinical benefit for people harboring any of these variants (Durmowicz AG+, 2018 [2]). As pointed out in the **Results**, the threshold of 10% functional improvement relative to WT function should not be considered as a rigid cut-off. Different statistical models and outlier removal could result in different calls around the threshold. For example, 10 variants approved by the FDA for modulator therapy fell under the 10% threshold (see above). An additional cautionary case is the N1303K variant, the fourth most common CF-causing CFTR variant based on the CFTR2 database. It is not currently approved for any modulator treatment, and with a functional improvement to 9.4% of WT CFTR following treatment with ELX/TEZ/IVA it did not surpass the 10% threshold. Yet clinical studies in several countries report that pwCF harboring the N1303K variant experience significant benefit from Trikafta® treatment. (Burgel+, 2023 [10] and Sadras+, 2023 [11]), consistent with these results.

An additional 224 variants with baseline function greater than 10% of WT also improved by more than 10% of WT function after drug treatment. While variants with higher baseline function often are associated with milder CF, of course depending on the severity of the variant on the other allele, there is no question that many people harboring these variants could still benefit from drug treatment, as demonstrated by clinical data for approved variants that fall into this functional range. Several variants with baseline function above 10% of WT, yet known to cause disease, were reported by Raraigh and colleagues (Raraigh KS+, 2018 [13]), and a recent case for the c.328G>C (p.Asp110His; legacy: D110H) variant was published by Jain and Cousar (Jain & Cousar, 2021 [14]).

Most of the 655 assessed variants had baseline functions <50% relative to WT CFTR, but there also were 104 variants with function >50%. This finding is not entirely surprising, given that DNA was collected in some countries based on symptoms as the only disease indicator (i.e., no confirmatory sweat chloride testing for some individuals). For 72 of these 104 variants, the underlying allele count is less than three and 61 of those have an allele count of one in the CFTR2 database. In addition, some of the variants with high baseline activity (i.e., forskolin response) in the functional assay may not show their pathogenicity in a cDNA-based assay format as other sequence variations not present in the experimental sequence may be required to elaborate the defect. Such CFTR variants might alter normal pre-mRNA splicing in the native gene context, or gene variants could severely disrupt CFTR-mediated bicarbonate transport (not captured in our functional assay), but not chloride transport. For example, in the cDNA context, the CFTR gene variant c.3700A>G results the predicted protein variant p.Ile1234Val (legacy: I1234V) that displays relatively normal CFTR function, 79.2% of WT function in our assay, yet this variant is reported as CF-causing in CFTR2. In fact, the variant creates a cryptic splice site that in the full gene context mostly results in a CFTR protein with six deleted amino acids, p.Ile1234_Arg1239del (a.k.a. I1234Vdel or I1234del6). This deletion variant has significantly reduced function (this study; Phuan PW+, 2021 [15]; Molinski SV+, 2014 [16]) and is likely the cause for CF in people harboring this variant. Other exonic variants with known or suspected splicing defects that were tested in this study include: c.454A>G (legacy: M152V), c.523A>G (legacy: I175V), c.1209G>C (legacy: E403D), c.1766G>A (legacy: S589N), c.2210C>T (legacy: S737F), c.2816A>G (legacy: H939R), c.2908G>C (legacy: G970R), c.2908G>A (legacy: G970S), c.3717G>C (legacy: R1239S), c.3872A>G (legacy: Q1291R), c.3873G>C (legacy: Q1291H) (Joynt AT+, 2020 [17], Raraigh KS+, 2022 [18]).

Some of the tested variants were originally observed in the context of complex variants, i.e., multiple in *cis* variants in a single allele. As expected, not all these variants are pathogenic when assessed by themselves nor can their responsiveness to the modulators be expected to always translate to other sequence backgrounds. Thus, the authors do not suggest that every person harboring a CFTR variant with >50% WT function (based on data from this study) should be treated with CFTR modulators. However, when people with such a variant present with clinical CF (presumably caused by other CF-causing variants), they may benefit from ELX/TEZ/IVA treatment. On the other hand, McCague and colleagues have shown that a cohort of pwCF with *CFTR* genotypes resulting in less than 2% of WT function sees their lung function measurement, measured as forced expiratory volume in 1 second relative to normal (FEV1% predicted), decline to below 60 by around 27 years of age, over 35 years earlier than pwCF with genotypes that confer 5-10% of WT CFTR function (McCague+ AJRCCM 2019 [19]). These data suggest the threshold for CFTR variant inclusion on the FDA drug label, drug-dependent improvement by 10 percentage points of WT CFTR function, is too strict and that pwCF with less than 5% of WT CFTR baseline function may experience meaningful clinical benefit from as little as a five percentage points improvement.

While disagreement in CFTR variant function and response to modulators between cDNA-based and native cell models was expected for a small number of variants, variant functional and biochemical data from this study are overall in good agreement with published data from CFTR2, Vertex (Yu H+, 2012 [20], Van Goor F+, 2014 [21]), and Cutting lab (Han ST+, 2018 [22]). For example, out of 110 variants annotated as CF-causing (CFTR2_7April2023 annotation) tested in this study, only five have an averaged baseline function of >30% of WT: c.1651G>A (p.Gly551Ser; legacy: G551S), c.3197G>A (p.Arg1066His; legacy: R1066H), c.3752G>A (p.Ser1251Asn; legacy: S1251N), c.3719T>G (p.Val1240Gly; legacy: V1240G) and I1234V, which as described above is not consistent with being a CF-causing variant when evaluated in a cDNA model. For G551S, R1066H, and S1251N, the forskolin response at UT is much higher than the combined average from the other two sites (which is <30% of WT). V1240G is borderline with an averaged baseline function of 34% of WT. Similarly, none of the 18 non-CF causing variants (CFTR2_7April2023 annotation) tested in this assay had a baseline CFTR function below 30% of WT. For correlation plots (using comparable datasets) between this study and published data from Vertex and Cutting group, see **supplemental Figures S4 and S5**.

Translation of modulator responsiveness from cDNA-based *in vitro* assays or patient-derived primary cell models into humans has been demonstrated for a good number of CFTR variants, from G551D and F508del to more rare variants (Yu H+, 2012 [20]; De Boeck K+, 2014 [23]; Van Goor F+, 2014 [21]; Burgel PR+, 2023 [11]), thereby providing further support for this approach. However, as pointed out above, cDNA-based models have limitations. For a select few variants these models do not fully represent the processing of the native CFTR gene with a given variant, e.g., variants I1234V, as detailed above, or G970R (Fidler MC+, 2021 [24]), and more broadly, the measured functional improvement for any given variant may be more indicative of a population level response rather than predictive for the response in a specific individual with CF. Even in the clinical trials for highly effective modulator combinations there were a few individuals that did not experience improvements in lung function (FEV1) or sweat chloride concentration (Keating D+, 2018 [25]; Davies JC+, 2018 [26]). For a detailed overview and discussion of preclinical model systems and personalized medicine in the context of CFTR modulator theratyping, see Clancy JP+, 2019 [27].

### Comparison with other in vitro data

Prior to this study, the largest and most important *in vitro* data set assessing modulator responsiveness of specific CFTR variants was generated by Vertex Pharmaceuticals and led to the approval of Trikafta® treatment for the 178 variants listed on the FDA drug label (FDA, 2021rev [3]). Of the 174 variants approved for Trikafta® therapy that were included in this study, for 164 (94%) the threshold for approval was reproduced, i.e., treatment-dependent CFTR function improvement was at least 10% of WT CFTR function. Of the ten variants that did not reach the threshold, five are within a 2.5% error margin below the 10% mark. Considering that the current data set was generated using a different assay, possibly with a different DNA base sequence, and across three different labs, this high level of alignment is impressive. In contrast to our study, Vertex generated the data in stably transfected FRT cell lines (as described in Van Goor F+, 2014 [21]) whereas this study employed transiently transfected FRT cells. Vertex used drug concentrations of 10 μM for ELX and TEZ (Fred Van Goor, personal communication) whereas 3 μM ELX and 3.5 μM TEZ were used in this study. Vertex produced all their data in one internal laboratory, whereas the data for this study was generated across three different laboratories and even more different operators. Overall, this excellent agreement between the results from Vertex and this study further increases confidence in the data that led to the approval of the variants on the FDA label, and equally it provides strong credibility for the measurements for variants that had not been assessed previously. Moreover, the coherence suggests that going forward the transient transfection method is of utility in very rapidly accessing new variants as they are discovered.

Stably transfected CFBE41o- or FRT cells were used in another previous study assessing 57 rare CFTR variants for responsiveness to ivacaftor treatment (Han ST+, 2018 [22]). The results from Han and colleagues and results generated in this study is in good agreement (**Figure S5**).

### Implications and use of the data from this study

The data from this study are not intended to enable FDA approval of additional CFTR variants for Trikafta® therapy. However, there is precedent of people with CF together with their CF care physician to petition their health insurances for off-label use of modulator therapy based on *in vitro* data. In most cases the *in vitro* data was generated with cells obtained from the person with CF, either through curettage or cytology brushing of the nasal mucosa of the inferior turbinate (Brewington JJ+, 2018 [28]; Pranke I+, 2019 [29]) or from rectal biopsy tissue (Berkers G+, 2019, [30]). There is a substantial cost-advantage of drug approval by variant, a precision medicine approach, versus personalized approval based on primary cell testing. A high correlation of *in vitro* to in vivo data has emerged in recent years, which together with strong drug safety data has enabled FDA-approval of Trikafta® therapy for many variants and ultimately many people with CF. It is the goal of this current study to assist additional people with CF, whose variants are so far not included on the FDA drug label, to petition for an off-label trial period of Trikafta® therapy and to demonstrate benefit based on established clinical readouts such as FEV1 and/or sweat chloride concentration, thereby justifying continued drug therapy. Furthermore, the data reported will hopefully inform the drug maker (Vertex Pharmaceuticals) regarding additional variants that may be Trikafta®-treatable and encourage steps for official FDA label inclusion. Many of the rare variants included in this study are predominantly found in people of color and other minoritized populations. Extending drug eligibility to additional CFTR variants will be a critical step in addressing the racial/ethnic inequities in CF care and outcomes.

## Acknowledgements

The authors acknowledge D. Benjamin for essential project management support, A. Vetter for the technical development of the WES western analysis, and C. Cotton for critical discussion and comments on the manuscript. This research would not have been possible without the data contributions of individuals with CF and the clinicians, data managers and analysts who provided these data to the Clinical and Functional TRanslation of CFTR (CFTR2) project.

## Funding

The work was supported by the Cystic Fibrosis Foundation [BRIDGE21XX1 to R.J. Bridges, THOMAS2018GO to P.J. Thomas, CUTT13A1 to G.R. Cutting and direct funding of the CFFT Lab]

## Author contributions

PJT, WRS, RJB, HB and MM conceived study and provided oversight. KSR and GRC provided DNA variant list and genetic expertise. LM, PB, ATP, JCh, VB, IM, and NEA carried out experiments. JCo, AL, NEA and AS carried out variant plasmid sequence analysis. JCo coded a raw data viewer, and JC, HB and AS collated data and performed data QC. NS, ASM and AS wrote code and performed statistical computing. AS described all the statistical analysis. HB and MM wrote the manuscript. WRS, RJB, ASM, KSR and GRC provided critical feedback on the manuscript.

## Declaration of Competing Interest

The authors declare no competing interests.

## Footnotes

1 The term ‘variant’ refers to a change in DNA that is an established change in DNA sequence that has been inherited and may or may not cause disease. The term ‘mutation’ describes the process of creating a DNA variant and it has adopted the implication of a negative effect in the public domain (PMID: 24387988). As we are studying DNA changes that are established, inherited and most are of unknown effect, the authors exclusively use “DNA variant” or “variant” throughout this report.

## Notes

### Competing Interest Statement

The authors have declared no competing interest.

