## Supplemental Figures and Methods for "*In Vitro* Modulator Responsiveness of 655 *CFTR* Variants Found in People With CF"

### Supplement: Tables and Figures

### Table S1

### Integrated Protein and Function data (site averages)

See **Table S1 upload - CFTR Variant Theratyping Bihler et al July-7-2023.xlsx**

Table S1: Functional and protein data from 655 CFTR variants and WT CFTR. Percent WT values for functional responses are given as across site averages. Variant functional responses (%WT) to IVA or ELX/TEZ/IVA are not explicitly reported here, but can be calculated from the reported values, e.g. variant response to IVA = Fsk+IVA (%WT) - Fsk+DMSO (%WT); variant response to ELX/TEZ/IVA = Fsk+IVA+(ELX/TEZ) (%WT) - Fsk+DMSO (%WT). For details regarding data generation, QC, analysis, and statistics, see the method sections in the manuscript body and supplement.

**Figure S1: Cross-site concordance of functional readouts**

Variant baseline function (%WT)

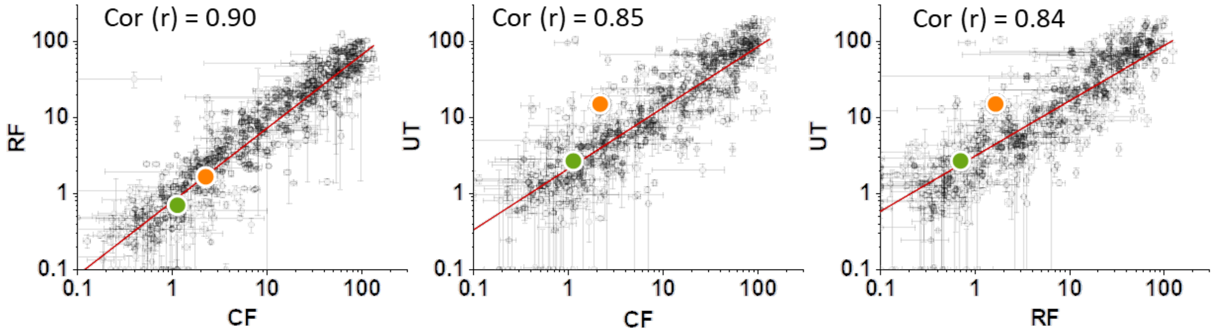


Variant response to **E**LX/**T**EZ/**I**VA (%WT)

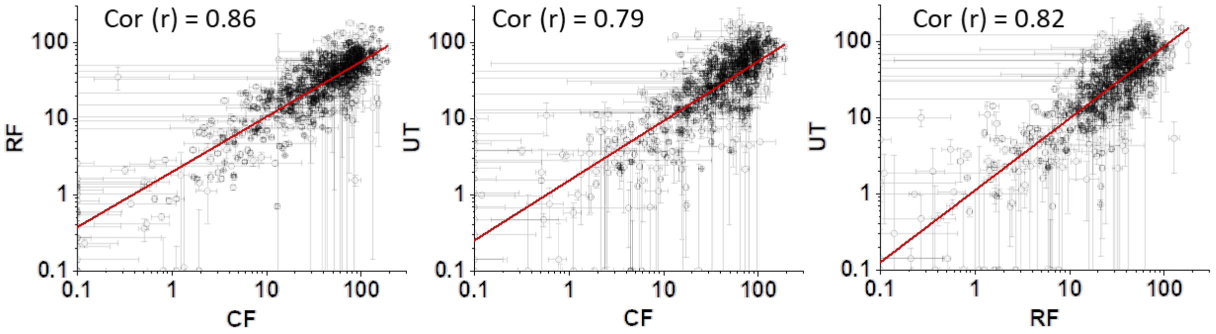


**Figure S1**: Cross-site concordance of functional readouts (mean ± SEM, for majority of variants, where n ≥ 1). Each graph compares the mean variant response, i. e. the CFTR mediated change in transepithelial conductance from the respective baseline (as percent of WT) from two sites for all matching variant samples. Total number of variant samples included for analysis equals 659 (=656 blinded samples plus 3 intraplate control samples). Red line represents best linear fit for correlation. **Top** panel: correlation plots and Pearson coefficients (r) for variant baseline function: ΔFsk (24h DMSO, Figure 1B). Overlap: CF/RF= 658, CF/UT = 611, RF/UT = 612 variant samples; Colored dots represent mean baseline function from assay plate controls for F508del (green) and G551D (orange). **Bottom** panel: correlation plots and Pearson coefficients (r) for variant responses to the triple combo: ΔELX/TEZ/IVA (cf. Figure 1B). Overlap: CF/RF = 655, CF/UT = 607, RF/UT = 606).

Correlation between CF and RF is stronger than the correlation between CF and UT or RF and UT as illustrated by the correlation coefficients and the orange dots (Figure S1, top panel), which represent the mean G551D baseline function from 205, 194, or 172 assay plates (from CF, RF, and UT sites, respectively). Technical reasons at UT, leading to an electrode contamination with IVA, may explain this discrepancy. As a consequence of electrode contamination with CFTR potentiator IVA, minute amounts of IVA in the top and bottom wells of UT assay plates prior to addition of forskolin may result in a somewhat increased baseline response of IVA responsive variants (e.g. G551D) upon addition of forskolin as well as in a reduced IVA response upon addition of IVA.

### Figure S2: Comparison of CFTR variant response to IVA and ELX/TEZ/IVA

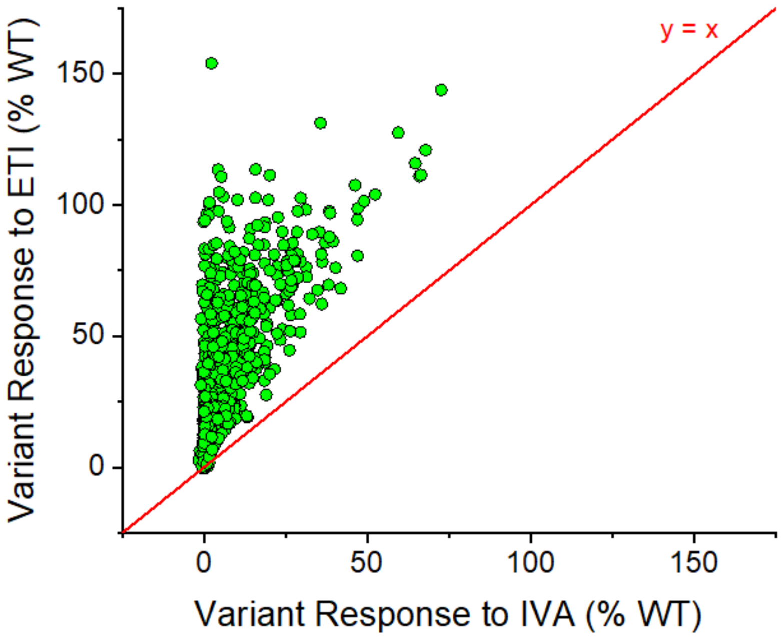


**Figure S2:** Functional Response of 656 CFTR Variants including WT to **E**LX/**T**EZ/**I**VA vs IVA relative to forskolin stimulated WT CFTR function. All variants had greater response to the triple combination **E**LX/**T**EZ/**I**VA than to ivacaftor (IVA) alone.

### Figure S3: Correlation between CFTR variant function (± IVA) and protein expression levels

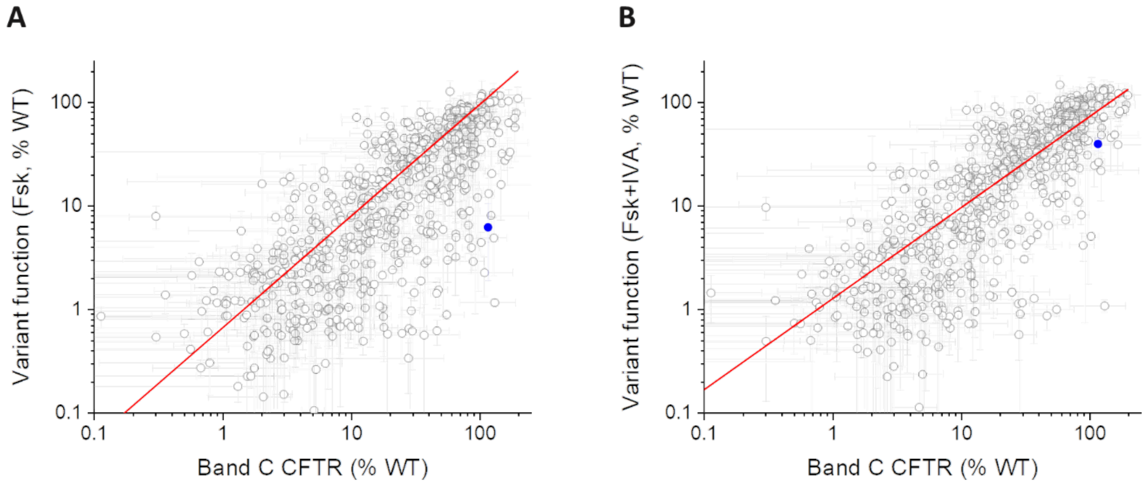


**Figure S3:** Correlation plot comparing CFTR Cl⁻ transport activities of 655 CFTR variants to mature variant CFTR protein expression levels (Band C, %WT). Red lines represent the line of best fit using a weighted linear regression. **A)** Comparison of forskolin stimulated (baseline) Cl⁻ transport activities to mature variant CFTR protein expression levels. Pearson's correlation coefficient (r) = 0.89 ($p = 2.7 \times 10^{-216}$), indicating a strong correlation between the two datasets. **B)** Comparison of forskolin plus ivacaftor (IVA) stimulated Cl⁻ transport activities to mature variant CFTR protein expression levels. Pearson's correlation coefficient (r) = 0.82 ($p = 4.3 \times 10^{-155}$), indicating a strong correlation between the two datasets. For reference, in-plate control G551D is highlighted in blue to demonstrate the effect of potentiator IVA on the class III variant G551D, which has a functional (gating) defect, but is expressed at normal WT CFTR levels.

**Figure S4: Variant protein expression – comparison with Van Goor et al, 2014**

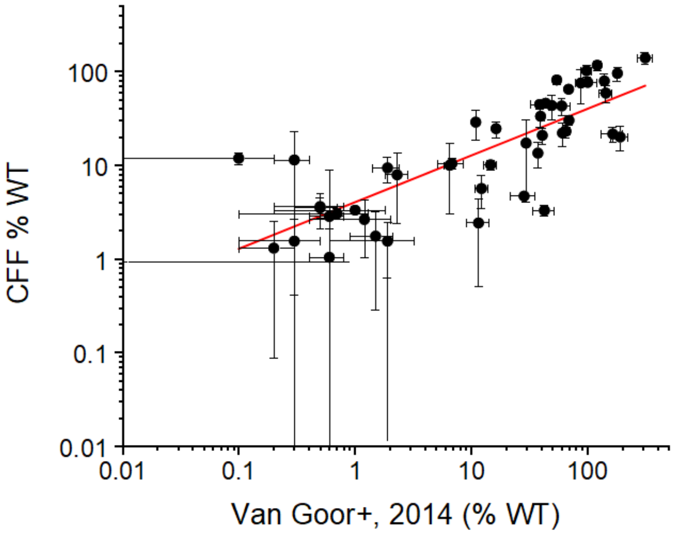


**Figure S4:** Correlation plot comparing levels of mature variant CFTR protein relative to mature WT CFTR protein (%WT) for a subset of 51 CFTR variants, either transiently (this study, CFF) or stably (Van Goor et al, 2014 [S1]) expressed in FRT cells. The red line represents the line of best fit using a simple linear regression analysis (no weighing) with a Pearson's correlation coefficient of (r) = 0.8 (p = 1.9 x $10^{-11}$), indicating a strong correlation between the two datasets.

### Figure S5: Variant baseline activity and IVA response – comparison with Raraigh et al., 2018 and Han et al, 2018

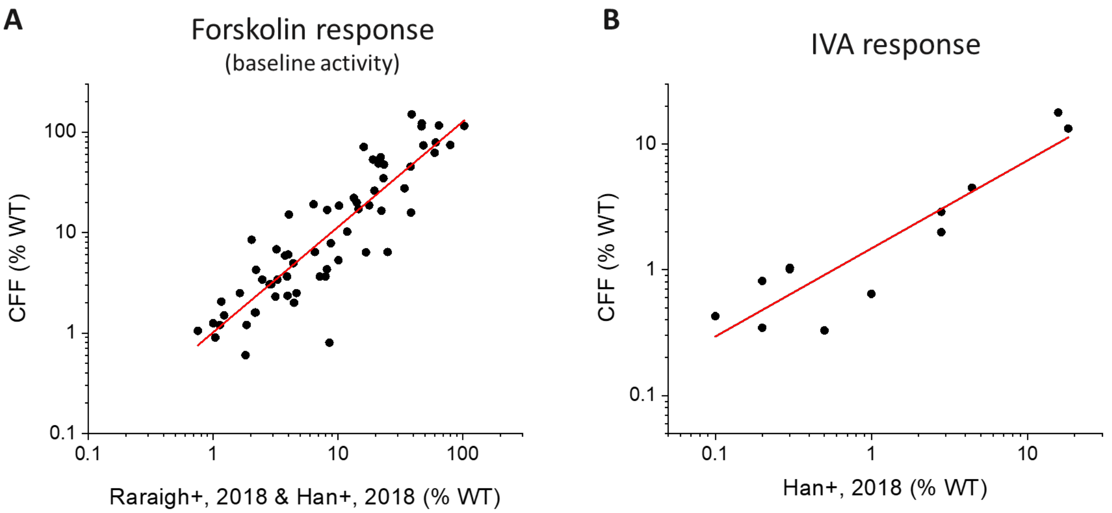


**Figure S5**: (**A**) Correlation plot comparing forskolin-stimulated (baseline) $Cl^-$ transport activities for a subset of 65 CFTR variants relative to WT CFTR baseline function (%WT). Variants were transiently expressed in FRT cells (this study, CFF) or stably expressed in CFBE cells (Raraigh et al, 2018 [S2]; Han et al, 2018 [S3]). The red line represents the line of best fit using a simple linear regression analysis (no weighing) with a Pearson's correlation coefficient of ® = 0.88 (p = 3.1 x $10^{-22}$), indicating a strong correlation between the two datasets. (**B**) Correlation plot comparing functional response of a subset of 18 CFTR variants to ivacaftor (IVA) as percentage of normal CFTR function (%WT). Variants were either transiently (this study, CFF) or stably (Han et al, 2018 [S3]) expressed in FRT cells. The red line represents the line of best fit using a simple linear regression analysis (no weighing) with a Pearson's correlation coefficient of (r) = 0.92 (p = 2.9 x $10^{-5}$), indicating a strong correlation between the two datasets.

### Figure S6: Variant surface expression - comparison with McKee et al., 2023

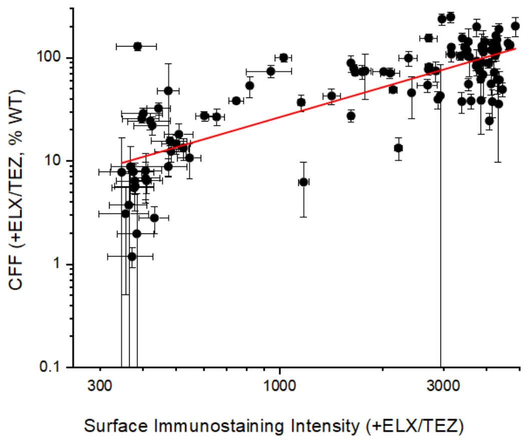


**Figure S6**: Correlation plot comparing levels of mature variant CFTR protein relative to mature WT-CFTR protein (%WT) from FRT cells (this study, CFF) and surface immunostaining intensity from HEK293 cells (McKee et al, 2023 [S4]) for a common set of 103 CFTR variants treated with CFTR modulators ELX/TEZ. The red line represents the line of best fit using a simple linear regression analysis (no weighing) with a Pearson's correlation coefficient of (r) = 0.79 (p = $3.6 \times 10^{-23}$), indicating a strong correlation between the two datasets.

**Supplemental Methods**

*Outlier detection and validation.* For each pDNA clone and treatment condition (vehicle or correctors), the three different sites were considered as groups and all measurements performed at the site (Fsk %WT or Fsk+IVA %WT) as a within-group sample – a classical one-way ANOVA design. It has to be noted that:

1. The number of observations per group is very small (two wells in most cases, with only a handful of larger samples due to repeated assay plates passing all the QC filters described in the text), leading to unreliable estimates of the variances.
2. Repeated measurements performed at the same site are distributed more tightly, consistently exhibiting smaller variance than the variance between the sites (*i.e.* between the per-site means). For instance, across all variants exhibiting average residual function FSK %WT in the 40-50 range, the mean and median of their standard errors of the mean (SEM) within each site are 5.2 and 3.8, respectively, while the mean and median of the SEMs of their averages across the sites are 11.3 and 10.9. For the variants with average residual function in the 10-20 %WT range, the corresponding average and median errors are 2.1 and 1.3 (within sites) vs 4.5 and 3.9 (between sites).

In order to address those issues, we start with the ANOVA model for the random variable representing repeated measurements of metric *r* (including treatment conditions, e.g. Fsk %WT with correctors) for plasmid *p*: $X_{pr} \sim \mu_{pr} + s + \varepsilon_{spr}$, where $\mu$ is the true underlying mean, *s* is the effect of different sites we aim to compare (categorical variable), and $\varepsilon$ is random variable representing measurement error. In standard ANOVA, the error (variance of the measurements) would be estimated for each model (i.e. each *p*, *r*) separately, from the small number of observations available for each variant. We postulate that there are no specific reasons for $\varepsilon$ to depend on a specific plasmid *p*, *except* for heteroscedasticity clearly observed in the data (variance increasing with the magnitude of the mean response). In other words, it is assumed that $\mathrm{Var}(\varepsilon_{spr}) = f(s, r, \mu_{spr})$, with no other specific dependency on *p*. Accordingly, the mean %WT responses $\hat{\mu}_{spr}$ estimated from the data at each site were stratified into tranches $T_i$ of width 10% and pooled variance values $\hat{\varepsilon}_{sri} = \bar{\varepsilon}_{spr}$: over $\hat{\mu}_{spr} \in T_i$ were calculated as averages of sample variances for given *s* and *r* across all variants in a given tranche. Those pooled values are expected to give more robust estimates of the true variance of the measurement error.

The one-way ANOVA calculation was implemented followed by post-hoc Tukey HSD comparisons between groups, using individual sample means of the responses at each site but pooled variances from the corresponding tranche in place of the variances estimated for each individual *p*. The described moderated statistic is not completely issue-free, still, because of (1) hierarchy of variances that is better modeled by mixed effect approach (see below), Var(labs) > Var(plates) > Var(repeats on a plate); and (2) effect of multiple testing, given that the comparison is performed independently for each of N=650+ variants x 2 treatment conditions x 2 response metrics (Fsk or Fsk+IVA %WT) = ~2600 ANOVA tests.

The calculated moderated ANOVA and Tukey pairwise HSD p-values still provide a useful and meaningful ordering of the variants by degree of (in)consistency among the measurements performed at individual sites. The heuristic approach used was as follows: for each *r* and *p* we order the per-site sample means $\hat{\mu}_{spr}$, let us denote the result $\{\mu_L, \mu_M, \mu_H\}$ (for low, medium, high); then the moderated post-hoc Tukey pairwise p-values $P_{ij}$ were examined, and $L$ was classified as a "low" outlier when *both* $P_{LM}, P_{LH} < 0.05$; similarly, $H$ was classified as the outlier on the "high" side when $P_{LH}, P_{MH} < 0.05$. Note that when only one outlier is detected according to this definition (either $L$ or $H$), the means at the other two sites ($MH$ or $LM$, respectively) are not significantly different ($P > 0.05$), *i.e.* the measurements at two sites out of three are consistent; when both $L$ and $H$ are classified as "outliers", this simply means that *all* sites are disagreeing with each other (this will be also the case when the data from only two sites are available and the measurements are inconsistent, $P_{LH} < 0.05$). The consistency counts obtained according to the above definitions are summarized in Table S3:

| | Fsk %WT | | Fsk+IVA %WT | |
|---|---|---|---|---|
| # outliers | DMSO | correctors | DMSO | correctors |
| 0 | 480 | 415 | 492 | 489 |
| 1 | 173 | 216 | 150 | 146 |
| 2 | 23 | 45 | 34 | 41 |

Table S2. The counts of cases of full consistency among the sites, 2 consistent out of 3, and fully inconsistent, per metric and per treatment. Note that the tests were run on the data for each pDNA clone; a few variants had multiple independently generated clones, thus the total count exceeds the count of distinct variants.

The moderated ANOVA p-values exhibit good correlation with post-hoc pairwise analysis (low p-value ≈ outlier(s) present) and provide a somewhat simpler way of viewing the data. The distribution across all pDNA clones suggests strong enrichment of small p-values, way above of what would be expected due to multiple testing alone (in which case the p-value distribution would have to be uniform). The enrichment must be due to a combination of truly inconsistent measurements and the hierarchical nature of the variance in the data, the latter not being properly accounted for in this simplified, ad-hoc approach.
The distributions of moderated ANOVA p-values for responses measured in the primary screen are shown in Figure S7. The figure also shows the distributions of the p-values for the plasmids selected for additional validation through assay re-runs (see main text). Note that the variants were selected for the assay repeat based on *any* response being a strong outlier (Fsk %WT or FSK+IVA %WT, vehicle- or corrector-treated). Multiple responses for the same variant, under different treatments might or might not have all shown up as outliers across the labs. Validation assays employed the same plate format as the one used in primary screen, with all treatments repeated in all cases, hence those responses of the selected plasmids that were consistent across the sites in the original screen can serve as negative controls for the follow-up validation experiment.

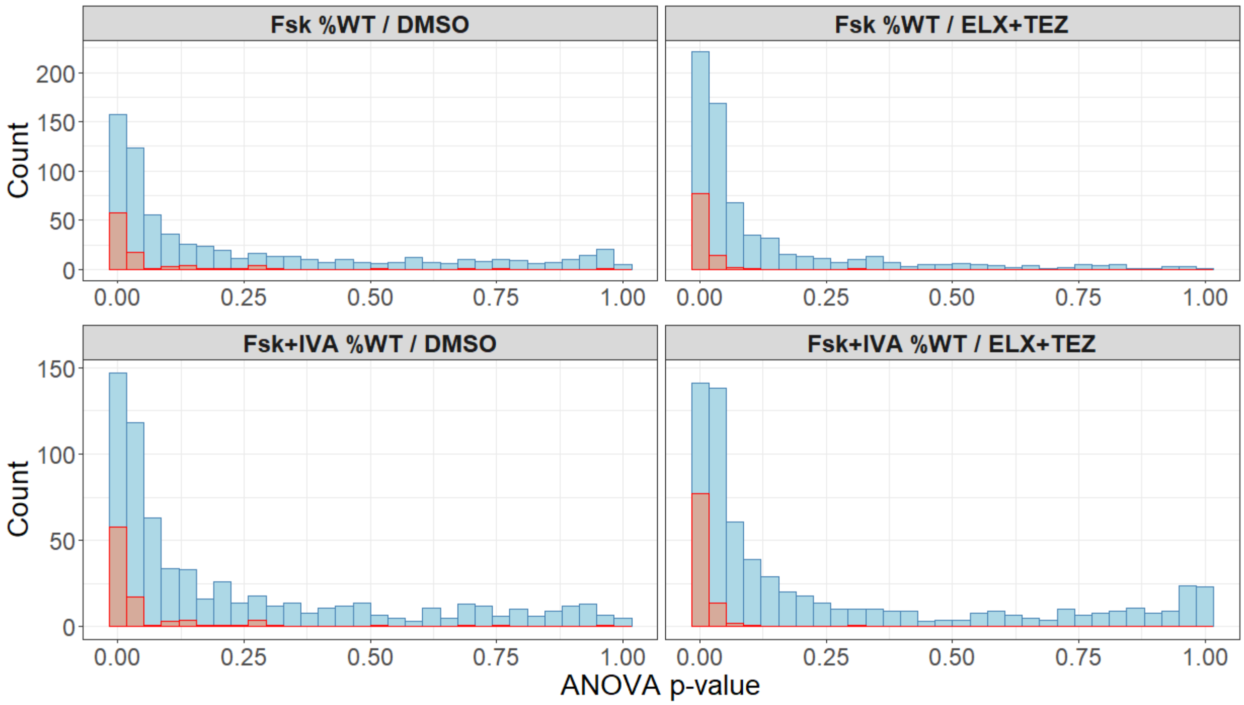


Figure S7. Blue: distributions of (moderated) one-way ANOVA p-values for the comparisons between samples of Fsk %WT and FSK+IVA %WT repeated measurements for each pDNA clone under vehicle or corrector treatments performed across the three sites. Red: distribution of the p-values of the 95 clones selected for follow-up validation assay.

Summary of validation assay results

| | | Primary assay | | |
|---|---|---|---|---|
| | | **no outliers** | **1 outlier** | **2 outliers** |
| **Validation assay** | **no outliers** | 77 | 116 | 11 |
| | **1 outlier** | 16 | 114 | 14 |
| | **2 outliers** | 2 | 6 | 24 |

Table S3: Contingency table of the consistency status of all the measurements (Fsk %WT, Fsk+IVA %WT, vehicle- and corrector-treated) across the sites for the 95 pDNA clones selected for validation. Primary: outlier status computed using primary assay data; validation: outlier status computed by replacing primary data with validation assay data for the site(s) where the assay was repeated.

The data show that out of 380 repeatedly assayed responses [95 pDNA x 2 (Fsk or Fsk+IVA %WT) x 2 (vehicle or correctors)], the consistency status across the sites did not change after the reassessment for 215 responses (57%), consistency improved for 141 (37%), and the number of outliers increased in 24 cases (6%). It is also worth noting that while the majority of the outlier calls stay unchanged, their significance as assessed by the pairwise post-hoc p-values does tend to decrease (p-values tend to increase, data not shown).

*Calculating average variant response*. As described above, the characteristic scales of the measurement errors (sample variances) have been observed to exhibit hierarchical structure, (wells on the same assay plate) < (different assay plates) < (different sites). Differences between plates, clones (when multiple pDNA clones are available for the same variant), or sites are random effects in this study. Hence, to compute the average response values for each variant under each treatment we employ the mixed effect model implementation `lmer` from the `lme4` R package [S5]. The most general model employed fits average value of each metric *r* (which represents here any of the metrics evaluated in this study, combined with either DMSO or corrector treatment) for variant *v* as random intercept-only mixed model $X_{vr} \sim (1|\text{assay plate}) + (1|\text{clone}) + (1|\text{site})$. Only the relevant terms would be used for fitting the average response of a particular variant when specific random effects are absent (*e.g.* there is a single assay plate from each site, or there's only a single pDNA clone available for the variant, or the data are coming from a single site only). Mean and standard error of the mean estimated from such mixed model are listed in Table S1.
